# Evolutionary tracking of cancer haplotypes at single-cell resolution

**DOI:** 10.1101/2021.06.04.447031

**Authors:** Marc J Williams, Tyler Funnell, Ciara H O’Flanagan, Andrew McPherson, Sohrab Salehi, Ignacio Vázquez-García, Farhia Kabeer, Hakwoo Lee, Tehmina Masud, Peter Eirew, Damian Yap, Beixi Wang, Jazmine Brimhall, Justina Biele, Jerome Ting, Sean Beatty, Daniel Lai, Jenifer Pham, Diljot Grewal, Douglas Abrams, Eliyahu Havasov, Samantha Leung, Viktoria Bojilova, Adam C Weiner, Nicole Rusk, Florian Uhlitz, Nicholas Ceglia, IMAXT consortium, Samuel Aparicio, Sohrab P. Shah

**Affiliations:** Computational Oncology, Department of Epidemiology and Biostatistics, Memorial Sloan Kettering Cancer Center, New York, NY 10065, USA; Tri-Institutional PhD Program in Computational Biology & Medicine, Weill Cornell Medicine, New York, NY 10065, USA; Department of Molecular Oncology, British Columbia Cancer Research Centre, Vancouver, BC, Canada; Irving Institute for Cancer Dynamics, Columbia University, New York, NY, 10027; Department of Pathology and Laboratory Medicine, University of British Columbia, Vancouver, British Columbia, Canada

## Abstract

Cancer genomes exhibit extensive chromosomal copy number changes and structural variation, yet how allele specific alterations drive cancer genome evolution remains unclear. Here, through application of a new computational approach we report allele specific copy number alterations in 11,097 single cell whole genomes from genetically engineered mammary epithelial cells and 21,852 cells from high grade serous ovarian and triple negative breast cancers. Resolving single cell copy number profiles to individual alleles uncovered genomic background distributions of gains, losses and loss of heterozygosity, yielding evidence of positive selection of specific chromosomal alterations. In addition specific genomic loci in maternal and paternal alleles were commonly found to be altered in parallel with convergent phenotypic transcriptional effects. Finally we show that haplotype specific alterations trace the cyclical etiology of high level amplifications and reveal clonal haplotype decomposition of complex structures. Together, our results illuminate how allele and haplotype specific alterations, here determined across thousands of single cell cancer genomes, impact the etiology and evolution of structural variations in human tumours.

## Introduction

More than 70% of human tumours are aneuploid and many harbor highly complex genomes^1^. Various processes including whole genome doubling^2^, whole chromosome and chromosome arm level gains and losses, segmental aneuploidies^1,3^, and complex structural rearrangements within and between chromosomes^4,5^ result in complex cancer haplotypes which can differentially impact maternal and paternal alleles. The degree or type of genomic instability correlates with clinical outcome in many settings, highlighting the importance of large scale genomic changes in interpreting therapeutic response^6–9^. Multi-region sequencing studies have begun to illuminate allele specific granularity of genomic instability^8,10^, yet how precise single cell-level variation of maternal and paternal alleles impacts genomic evolution remains understudied.

Recent advances in scalable low pass single cell (or nucleus) whole genome sequencing can profile large numbers of cells per sample (100’s to 1000’s) and offer new opportunities to define properties of intra-tumour heterogeneity in genomically unstable tumours^11,12^. Methods such as direct library preparation+ (DLP+) provide granularity to identify ongoing instability at high resolution^11^. Yet, most single cell whole genome profiling has focused on the analysis of total copy number due to technical limitations. Bespoke library preparation methods such as strand-seq^13^ can infer allele specific copy number but do not scale well to large numbers of cells, while high throughput methods require dedicated computational solutions due to their sparse coverage^14,15^. Total copy number approaches accordingly miss important events such as copy neutral loss of heterozygosity (cnLOH), impacting key biological attributes such as bi-allelic inactivation of tumour suppressor genes and mechanisms of immune evasion^16^. Furthermore, accurate decomposition of complex genomic variants requires mapping events to homologous chromosomes^17^.

We developed a new analytical method to identify allele and haplotype specific copy number in scDNA and applied it to DLP+ single cell whole genome sequencing of a cohort of more than 32,000 cells from 22 genomically unstable tumours, 7 genetically engineered cell lines and 3 patient derived cell lines. We used cell-level allele and haplotype specific CNAs to compute accurate phylogenetic trees and measure rates of instability, integrated them with single cell RNA sequencing to reveal properties of convergent copy number evolution and combined them with subclonal structural variants to infer sequential evolution of complex genomic changes. Our results highlight the prevalence of continual accrual of large genomic alterations, providing new insight into copy number driven evolution of cancer genomes at haplotype specific resolution.

## Results

### Accurate allele-specific copy number in single cells

To study the impact of allele and haplotype specific copy number alterations in single cells at scale, we developed an analytical approach called `schnapps` (single cell haplotype copy number analysis by phased probabilistic states). `schnapps` phases haplotype blocks across cells, computing a value for the B allele frequency (BAF) in 500kb bins across the genome. Phasing of alleles is refined based on global imbalances in clusters of cells sharing similar copy number events and allele specific states per bin are inferred using a hidden Markov model (HMM) that incorporates total copy number based on relative read depth and BAF’s (see methods).

We evaluated `schnapps` performance metrics on previously published single cell data from a group of ovarian cancer cell lines derived from the same patient^11,18^. This data includes cell lines derived from the primary tumour (SA1090), and 2 relapse specimens from the primary site (SA921) and ascites (SA922) respectively. Mean coverage per cell was 0.16X. We found clear examples of loss of heterozygosity (BAF = 0.0) at chromosomes 2p, 3p, 4p, 13, 16,17, 21 and 22 in individual cells (see **Figure 1a** for an example). BAF’s were distributed around the expected values, even at relatively high copy states (>8) (**Figure 1b**). We then computed the variant allele fraction (VAF) of clonal single nucleotide variants (SNV) per allele specific state. The VAF followed expected distributions, whereby mutations in balanced regions of the genome had VAF ∼ 0.5, mutations in homozygous regions had VAF ∼ 1.0, and mutations in unbalanced regions exhibited modes consistent with mutation acquisition timing pre and post the copy number alteration (e.g. 1/3 and 2/3 for 2|1) (**Figure 1c**).

**Figure 1.**
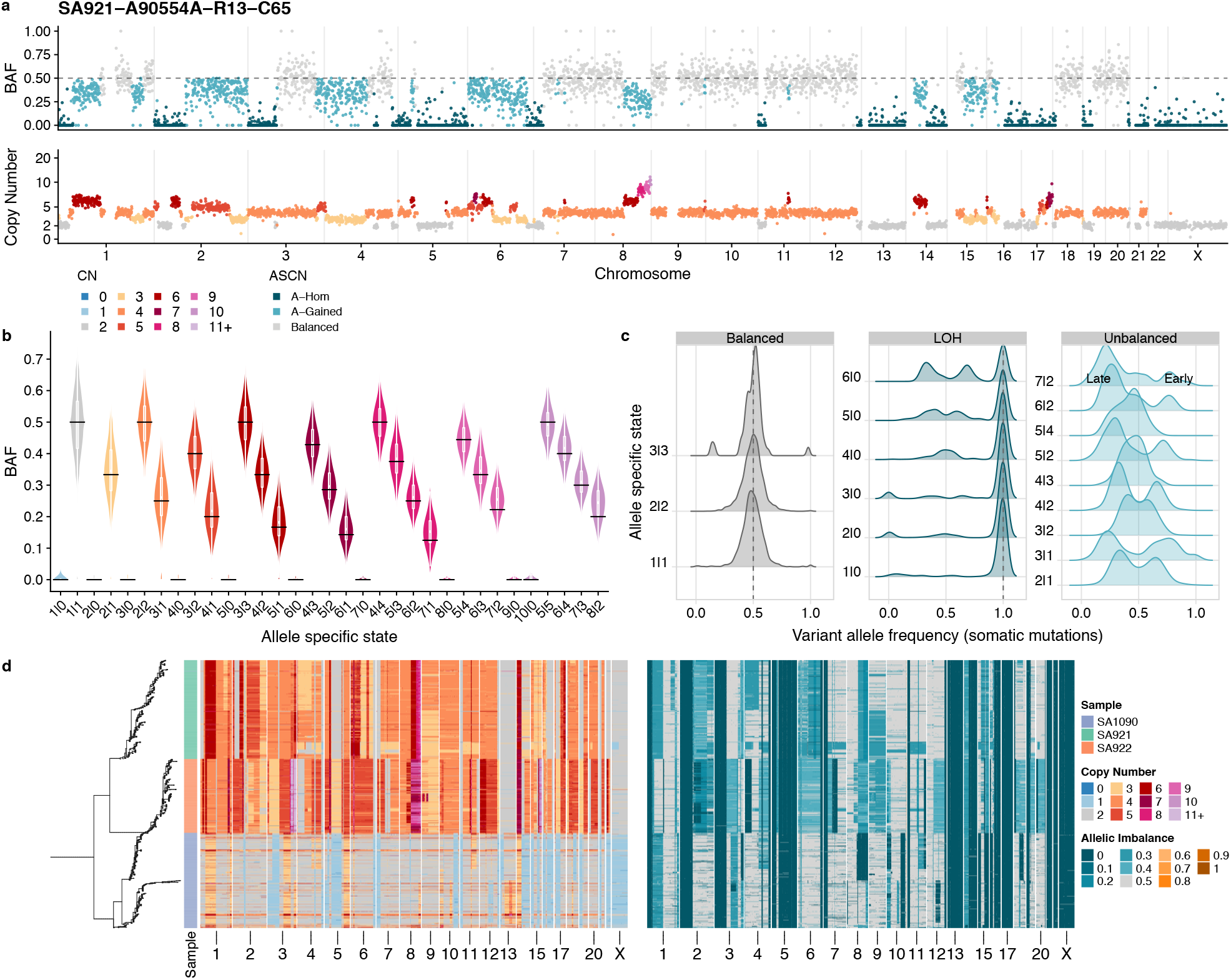
Allele specific inference at single cell resolution. a) Example allele specific copy number in a single cell. Genome position in 0.5Mb bins is shown across the x-axis, top panel shows the B-Allele Frequency (BAF) per bin coloured by inferred allele specific state, bottom shows the corrected read counts per bin colored by inferred total copy number. b) Distribution of BAF as a function of allele specific state across all cells in sample 2295. c) Variant allele frequency of somatic mutations mapped to allele specific states grouped by Balanced states (A==B), LOH states (A or B = 0) and Unbalanced (A!=B). d) Copy number heatmaps of all 1031 cells in OV2295, ordered by phylogeny computed from allele specific states per cell (left). Left heatmap shows total copy number, right heatmap is allele specific copy number.

Having confirmed the accuracy of allele specific inference we then computed a phylogenetic tree of all 1031 cells in this three sample dataset with a phylogenetic inference method, `sitka`^19^ using allele specific copy number as input (see methods). Visualizing this tree together with phylogenetically ordered heatmaps of total copy number and allele specific copy number revealed genomic alterations and clonal relationships that were not predicted with total copy number, **Figure 1d**. Firstly, we did not observe any total copy number events that were shared between the three samples; allele specific copy number however revealed that all cells are homozygous (BAF = 0.0) at chromosomes 5, 13, 17, 21 and 22, **Figure 1d**. In addition, allele specific copy number events such as cnLOH on chromosomes 8q and 10p in a subset of cells in SA1090 further refine clonally expanded populations, **Figure 1d**. Taken together, this analysis illustrates the increased resolution provided by allele specific copy number at the single cell level.

We next inferred allele specific copy number in a set 7 genetically engineered cell lines and 22 human tumours with DLP+ single cell whole genome sequencing^20^. The cell lines originate from a WT immortalized hTERT mammary epithelial cell line, from which we generated derivative lines, using CRISPR to inactivate key DNA repair pathway genes. Our data included: wild-type (*WT*, n=1), *TP53^−/−^* (n=2), *TP53^−/−^ BRCA1^−/−^* (n=1), *TP53^−/−^ BRCA2^−/−^* (n=2), and *TP53^−/−^ BRCA2^+/−^* (n=1). The human tumour data derives from 15 high grade serous ovarian cancer (HGSOC) samples and 7 breast cancer samples from both PDX models (n=19) and primary human tissue (n=3). A full description of each sample is provided in **Supplementary Table 1**. Allele specific copy number inferred from matched bulk whole genome sequencing were highly similar to the average allele specific copy number of single cells (**Supplementary Figure 1**). This, together with SNV VAF distributions confirmed the accuracy of our inferences for downstream analysis.

### Background copy number alteration and positive selection in 22 tumours

We next investigated the landscape of copy number alterations in the tumour cohort from single cell whole genome sequencing (median 697 cells per sample, range 49-2,627), enabling unambiguous cancer cell fraction (CCF) estimates in each tumour. We classified regions of the genome in each cell as LOH and gained or lost relative to cell ploidy, and then calculated CCF for each type of event across the genome. At the clonal level (CCF > 95%) recurrent events across tumours included 1q, 3q and 8q gains and prominent focal alterations around the oncogenes KRAS, MYC, CCNE1 and PIK3CA. Recurrent losses included 1p, 8p, 5p, 17p and 4, **Figure 2a**. These are all known recurrent events in breast and/or ovarian cancers, corroborated by the Pan-Cancer Analysis of Whole Genomes (PCAWG) cohort^5^, **Figure 2a**. In samples containing *TP53* and/or *BRCA1* loss of function mutations, invariably 100% of cells were homozygous around these loci, **Supplementary Figure 2**.

**Figure 2.**
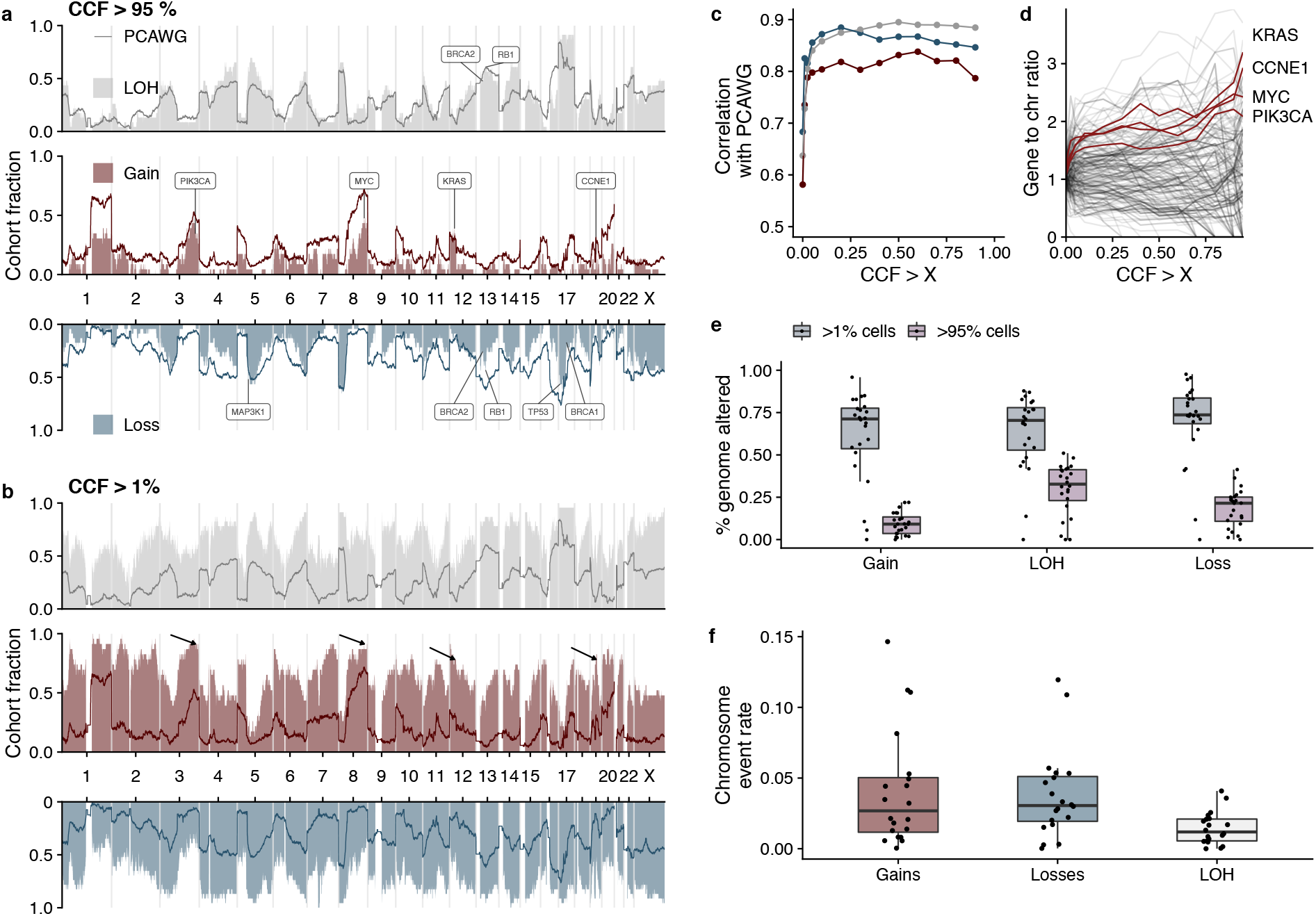
Landscape of copy number alterations as a function of clonality. a) Landscape of alterations (LOH, Gains and Losses) across the genome in 25 tumours. The fraction of tumours with a particular type of alteration is shown on the y-axis, position along the genome is shown on the x axis. Shown here are recurrent clonal alterations (CCF > 95% per tumour) and b) recurrent alterations with CCF > 1% per tumour. Darker colored lines show the PCAWG cohort frequency. Arrows on the gained track show the location of the oncogenes PIK3CA, MYC, KRAS and CCNE1. c) Correlation between frequency distributions and PCAWG frequency as a function of CCF. d) Ratio of gene frequencies to chromosome frequencies as a function of CCF. Trajectories of MYC, CCNE1, PIK3CA and KRAS are highlighted with red lines e) Fraction of genome altered by gains, losses or LOH for alterations present in 95% of cells and 1% of cells f) Chromosome event rate for gains and losses and LOH. Each data point represents the average chromosomal rate per sample.

In contrast to clonal alterations, recurrent alterations present in at least 1% of cells were more uniformly distributed across the genome (**Figure 2b**). To quantify these observations, we compared CNA distributions with those reported in the PCAWG cohort^5^. Correlation with the PCAWG data became stronger for losses, gains and regions of LOH as CCF increased (**Figure 2c**), and focal amplifications of key oncogenes became more pronounced (**Figure 2d**). Furthermore, a higher fraction of the genome was altered at CCF > 1% than CCF > 95% (**Figure 2e**). Chromosomal event rates (see methods) were estimated to be on average 0.01 per division per chromosome for LOH events (excluding losses resulting in 1 copy) and 0.03 for gains and losses (**Figure 2f**). This analysis revealed a high but consistently uniform background rate of copy number alteration across the genome, suggesting that chromosomes continually acquire CNAs without enrichment for specific genomic loci, and focuses attention on regions likely under positive selection in the initial clonal expansions of these tumours. Importantly, these regions include known oncogenes such as *KRAS*, *CCNE1*, *PIK3CA* and *MYC*.

### Parallel gains and losses inferred from haplotype specific copy number

We next leveraged our ability to phase alleles to individual haplotypes in single cells and estimate “haplotype specific copy number” to investigate parallel copy number evolution^8,10,14,15^ (subclones with the same total copy number but different haplotype configurations) as a putative signature of selection (**Figure 3a,b**). Parallel evolutionary events are often considered indicators of positive selection, reflecting convergence on a particular advantageous genotype^21,22^. We detected a striking example consistent with this interpretation in our engineered cell line data, where gain of chromosome 20 of both the paternal and maternal allele is observed in 100% (7/7) of these lines, often at high frequency (**Figure 3c**). This is more common for chromosome 20 than other chromosomes suggesting it provides a fitness advantage for cells in this system (**Supplementary Figure 3**).

**Figure 3.**
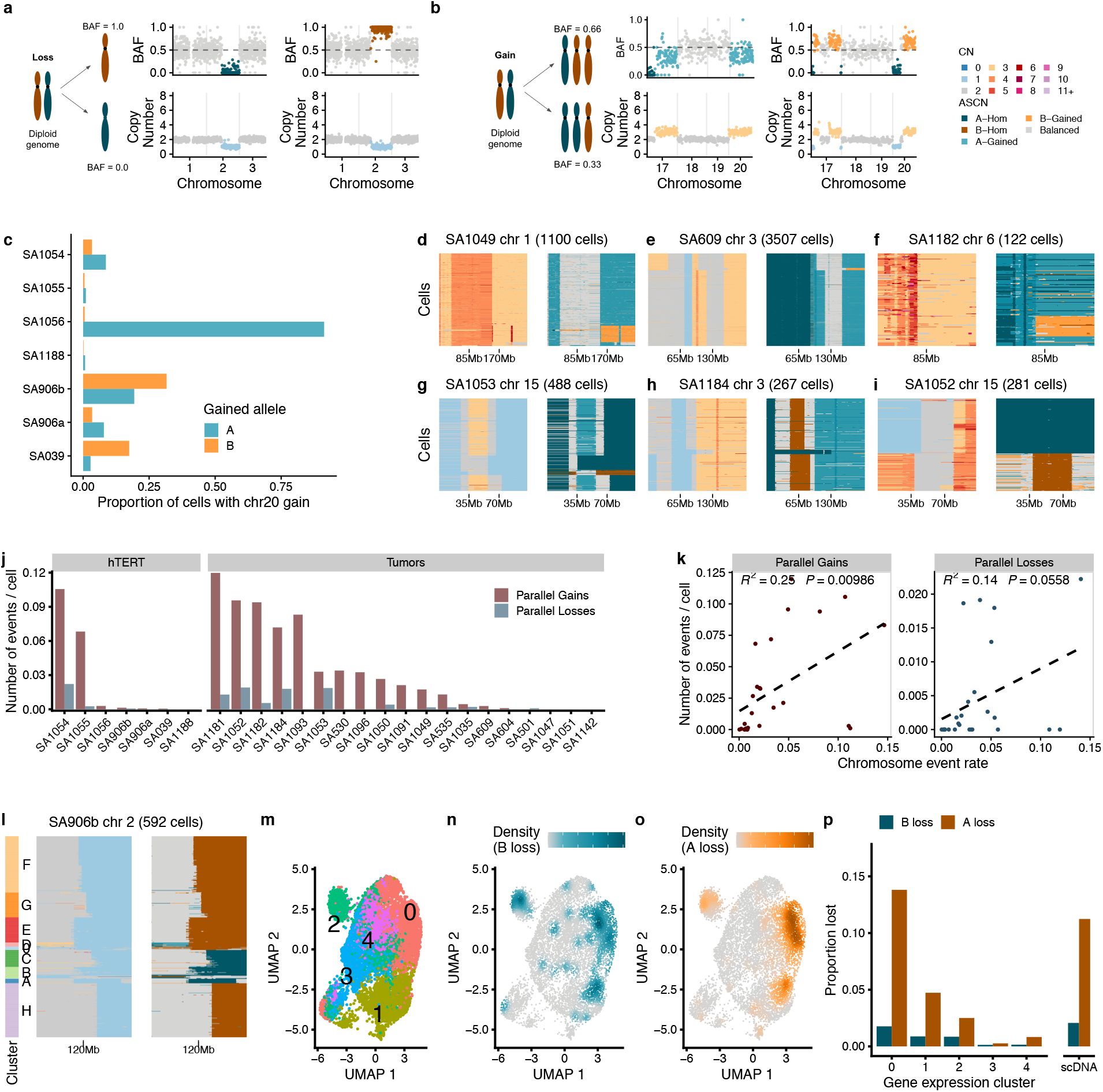
Parallel copy number evolution. a) Parallel losses at chr 2 in 2 single cells from SA906b b) Parallel gain in 2 single cells from SA906b showing gains in chr17 and chr 20 c) Proportion of cells with gains of chr20 in each allele in the engineered hTERT cell lines d)-i) Examples of parallel gains and losses in our data. Each heatmap shows a single chromosome or chromosome arm from a single sample, total copy number is shown on the left and the allelic imbalance on the right, colour coding the same as in panel b). Cells are ordered from top to bottom according to computed phylogenetic trees. j) Number of parallel events in each sample divided by the number of cells. k) Number of parallel events vs chromosome event rates e) Example of a parallel loss in chr2q in SA906b (TP53−/−), 592 cells are shown. Left heatmap shows total copy number, right heatmap shows allele imbalance. Distinct clusters of cells are labelled on the left. l) UMAP of gene expression of SA906b coloured by gene expression cluster. n) UMAP coloured by density of cells with the B allele lost and o) the A allele lost p) Proportion of cells with allele lost in each gene expression cluster (A lost = BAF < 0.1, B lost = BAF > 0.9) and in scDNAseq.

We then looked for parallel evolutionary events in the tumour samples. Examples of parallel gains include chr1q in SA1049 (**Figure 3d**), chr3q in SA609 (**Figure 3e**) and chr6q in SA1182 (**Figure 3f**) and parallel losses or cnLOH at chr15 in SA1053 (**Figure 3g**), chr3 in SA1184 (**Figure 3h**) and chr15 in SA1052 (**Figure 3i**). Even when observed on the same haplotype, many of the events had distinct breakpoints, consistent with these alterations occurring numerous times during tumour evolution. Overall, 18/21 tumours (3 tumours were removed from this analysis due to low cell numbers) and 7/7 cell lines had at least 1 parallel event present in more than 1% of cells, with parallel gains typically more common than losses (**Figure 3j**). Although some of these parallel events affected chromosomes commonly altered in HGSOC or TNBC, such as gain of 1q in SA1049, 3q in SA609 and chr15 losses, many were observed in chromosomes that are not recurrently seen in large cohorts^5^. This raises the possibility that parallel events may often arise by chance rather than due to selection of a particularly advantageous karyotype. Consistent with this, we found that the number of parallel events per tumour was positively correlated with chromosomal event rates (**Figure 3k**).

We next sought to explore whether parallel events produce convergent effects on transcriptional phenotype. In order to isolate the effect of a particular parallel event on transcription, we first identified a group of 592 cells in SA906b (TP53−/−) that had highly consistent copy number alterations apart from parallel losses on chromosome 2q (**Figure 3l, Supplementary Figure 4**). In order to assess transcriptional phenotype, we generated single cell RNA sequencing using the 10X platform and genotyped haplotype blocks identified in the scDNA in the scRNA profiled cells (see methods). Using the per cell counts we computed BAF values per chromosome arm and confirmed that this approach could accurately recover allelic imbalance in single cell transcriptomes (**Supplementary Figure 5**). We then clustered cells using Louvain clustering based on gene expression (**Figure 3m**) and identified losses of chr2q (BAF < 0.1 for loss of B, BAF > 0.9 for loss of A) in each cell. Gene expression clusters 0 and 1 were enriched for both types of losses (proportions test, p<0.001) confirming that this parallel copy number event results in a convergent effect on the transcriptome (**Figure 3 n,o,p**).

### Decomposition of complex structural rearrangements at haplotype resolution

Another striking source of variation between cancer cell genomes with functional consequences for tumour cell fitness is the variation in the level of oncogene amplification between cells^20^. A plausible mechanistic explanation for this is genome diversification through breakage fusion bridge cycles (BFBC)^23^, a known mechanism of complex rearrangements that can lead to amplifications of oncogenes^24,25^. We hypothesized that BFBC-like processes may generate diversity in the magnitude of oncogene amplification between cells and that progressive BFBC evolution could be resolved with haplotype specific copy number analysis.

BFBC typically result in reciprocal patterns of gains and losses in daughter cells following aberrant missegregation of chromosomes during cell division^17^ (**Figure 4a**). We identified this distinctive pattern in a subset of cells on chr 3 in SA1188 (*TP53*^−/−^ *BRCA2*^+/−^). Mapping events to homologous chromosomes revealed clusters of cells consistent with different stages of BFBCs (**Figure 4a,b**). Clusters B and F are consistent with the expectation of daughter cells following an initial BFBC, with a reciprocal gain and loss at the terminal end of chromosome 3 (**Figure 4a-d**). To further refine this analysis, we identified rearrangement breakpoints in these cells using pseudobulk analysis (see methods). These rearrangements further refined the likely BFBC progression. We can deduce that in cluster B a genomic segment at the end of homolog B first underwent a number of inversions and the “new end” fused with its sister chromatid generating a foldback inversion (FBI) (**Figure 4c**). FBIs - defined by genomic segments stitched together head to head - are another footprint of BFBC^26^. We also observed clusters of cells consistent with a second cycle producing either internal amplifications (clusters C, D, H, I and G) or extending the terminal loss (cluster E) on the same homolog (**Figure 4-g).** In addition, we identified a minor subpopulation of cells (n=19) with a focal amplification (total copy number = 5) around *PIK3CA* (**Figure 4f**), suggesting further BFBC cycles driving amplification of this oncogene. Independent phylogenetic reconstruction was consistent with the expected branching process induced by BFBC (**Supplementary Figure 6**). Other examples of BFBC mediated genomic variation in the cell lines included *MYC* amplification in SA906a (*TP53*^−/−^) and chr20 amplification in SA906b (*TP53*^−/−^) (**Supplementary Figure 7**).

**Figure 4.**
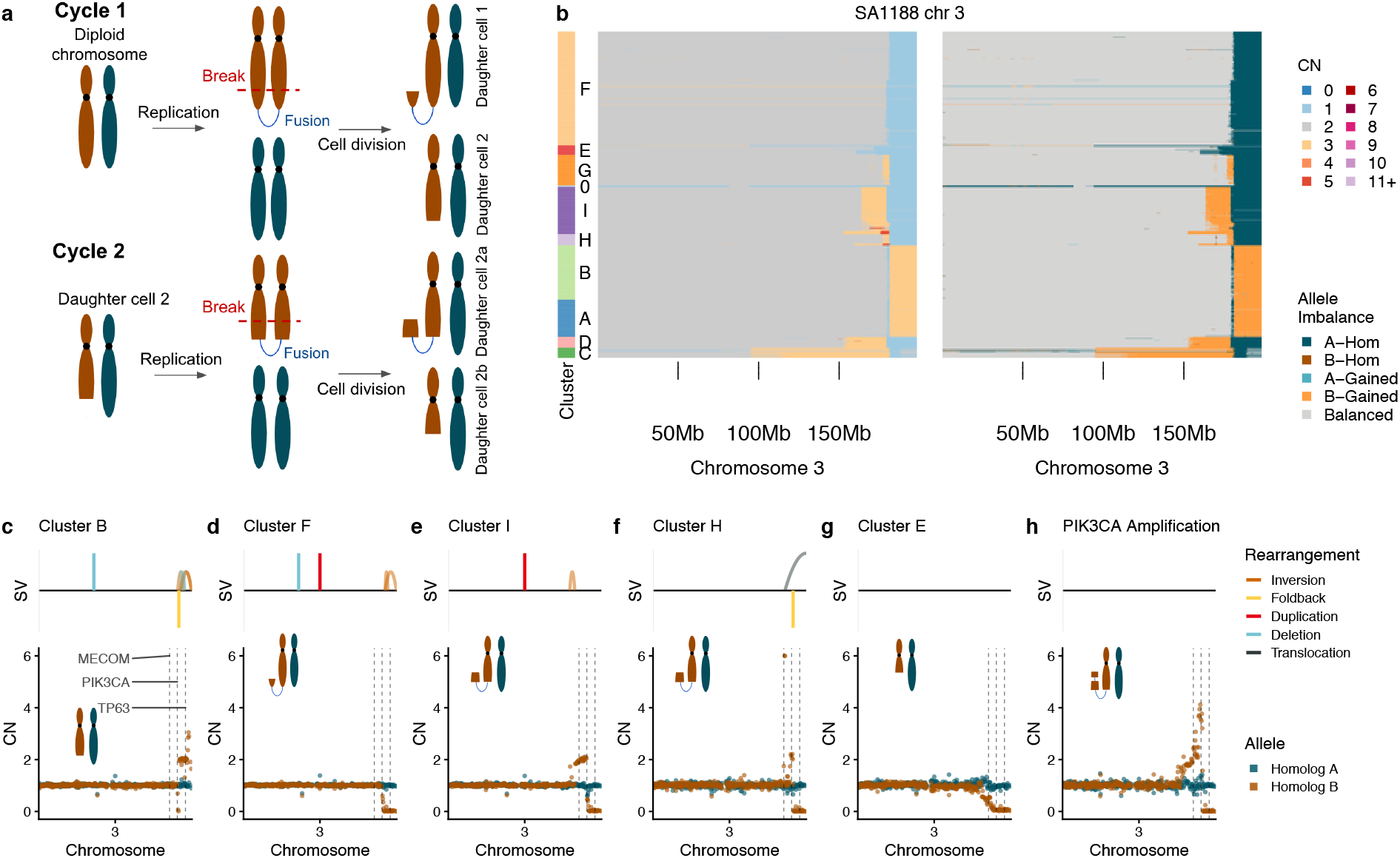
Breakage fusion bridge cycles in an *TP53*^−/−^ *BRCA2^+/−^* cell line. a) Diagram of breakage fusion bridge cycles b) Heatmaps of total copy number and haplotype specific copy number in SA1188. c)-h) Haplotype specific copy number and structural variation in clusters B,F, I, H, E and the small subpopulation with PIK3CA amplification. Here we plot the copy number for each homologous chromosome in brown for homolog B and blue for homolog A.

### BFBC results in diverse oncogenic amplifications in tumours over time

We next looked for the signature of BFBC in our tumours, focusing on 3 key characteristics: i) amplifications adjacent to a loss ii) identification of FBI rearrangements and iii) a ‘staircase’ pattern of copy number alterations (**Supplementary Table 2**). In order to dissect the relative timing of BFBC’s, we first focused on 2 PDX samples (SA1035 & SA535) that were serially passaged over time. In SA1035, there were numerous subpopulations in chr19 consistent with a simple BFBC. Evidence for BFBC included internal amplifications adjacent to a terminal loss on the same homolog and congruent FBI breakpoints (**Figure 5a**). One of the larger subpopulations (cluster C), included an amplification of *CCNE1* (total copy number 4) (**Figure 5a**). In SA535 we observed a more complex BFBC-like rearrangement pattern encompassing the *FGFR1* locus, with distinct amplitude differences between subpopulations (ranging from CN=2 to CN>8). Rearrangement breakpoints in cluster C suggest that here, the chromosome was stabilized via fusion with chromosome 5, while in cluster E the chromosome was stabilized via a complex rearrangement involving both arms of chromosome 8 **Figure 5b**. Investigating single copy number profiles revealed rare cells where the amplification was completely absent, cells where there was a small amplification (copy number = 3) and cells with extreme copy number (copy number > 10) (**Supplementary Figure 8**). We then computed the frequency of each cluster at each timepoint in both SA535 and SA1035 and found that all clusters were represented at a non zero fraction (albeit often at very small frequencies) in the first time point, **Figure 5c**. In SA535 the population of cells with *FGFR1* copy number < 3 (cluster W) remained at a low frequency over time, while in SA1035 the *CCNE1* amplified subclone (Cluster B) clonally expanded and became the dominant subclone by passage 8.

**Figure 5.**
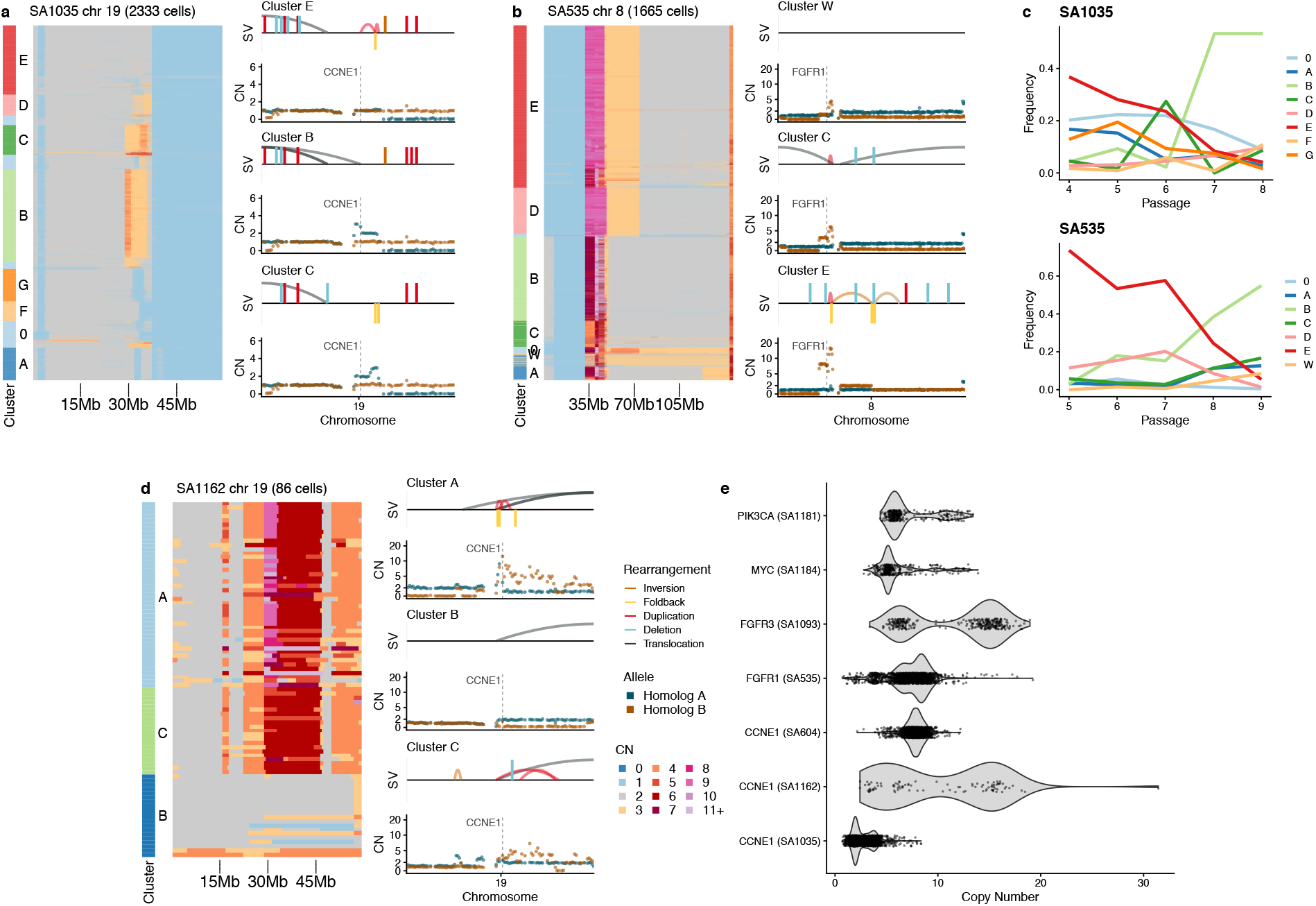
BFBC in human tumours. a) BFBC in chr19 SA1035. Left shows heatmap of total copy number for 2333 cells in chromosome 19 clustered using UMAP and HDBSCAN. Right hand side shows pseudobulk average haplotype specific copy number profiles of 3 clusters with distinct events. Schnapps is used to infer the copy number in the 2 homologous chromosomes and these are plotted together with brown points indicating allele B and blue points indicating allele A. Top track for each copy number profile shows the structural variants found in these clusters. b) and d) are equivalent to a) for chr8 in SA535 b) and chr19 in SA1162 d). c) Frequency of each cluster across time in SA1035 and SA535 e) Distribution of the raw copy number per cell in cases that were consistent with BFBC induced oncogene amplification.

We also observed BFBC driving subclonal amplification of *CCNE1* in SA1162, one of the primary human tumour samples. In this patient we observed subclones with 2, 4 and >15 copies of *CCNE1* **Figure 5d**. Other examples that were consistent with oncogene amplification due to BFBC include *KRAS* in SA604, *MYC* in SA1184, *FGFR3* in SA1096 and *PIK3CA* in SA1181, **Supplementary Figure 9**. In all these cases we observed considerable genomic variation between cells leading to variable levels of oncogene amplification **Figure 5e**. Notably, we found that many of these genes had focal amplifications enriched at high CCF across the whole cohort, **Figure 2c**. Our data also revealed numerous occurrences of BFBC mediated genomic diversity that did not appear to be associated with oncogenic amplification, underlining that this process likely occurs in the background throughout tumour evolution, **Supplementary Figure 10**.

Other genomic instability processes implicated in oncogene amplification include chromoplexy^27^, ecDNA^28^ and tyfonas^25^. These processes often result in highly complex structural rearrangements across multiple chromosomes and in some instances, amplify several oncogenes simultaneously. We identified these types of events in multiple samples. In SA1049 we identified a complex event that included genomic segments from chromosomes 6,7,8,12, 19 and 20 and included amplifications of both *KRAS* and *FGFR1*, **Figure 6a**. Striking differences between cell clusters were apparent including co-amplification of genomic segments in chromosomes 6 and 12 resulting in a high level amplification of *KRAS* in cluster E. SA604 harbored a complex event that included chromosomes 6, 8, 12, 19 and 20 with amplification of *MYC*, *KRAS* and *CCNE1* **Figure 6b**. Again, variability in complex structural alterations between clusters was notable, including rearrangements between chromosomes 6 and 20 amplifying *KRAS* **Figure 6c**. Mapping the clusters identified in SA604 to passages revealed that all populations were present at the first time point and the cluster with low KRAS copy number (cluster A) remained at low frequency. Many of these inter-chromosomal amplifications also had clustered FBI’s and segments with variable copy number, suggesting that BFBC-type processes may contribute to the generation of these types of events^17^. Together these results reveal extensive variation in complex rearrangements as an underappreciated source of variation in cancer genomes that is often obscured in bulk sequencing of tumours.

**Figure 6.**
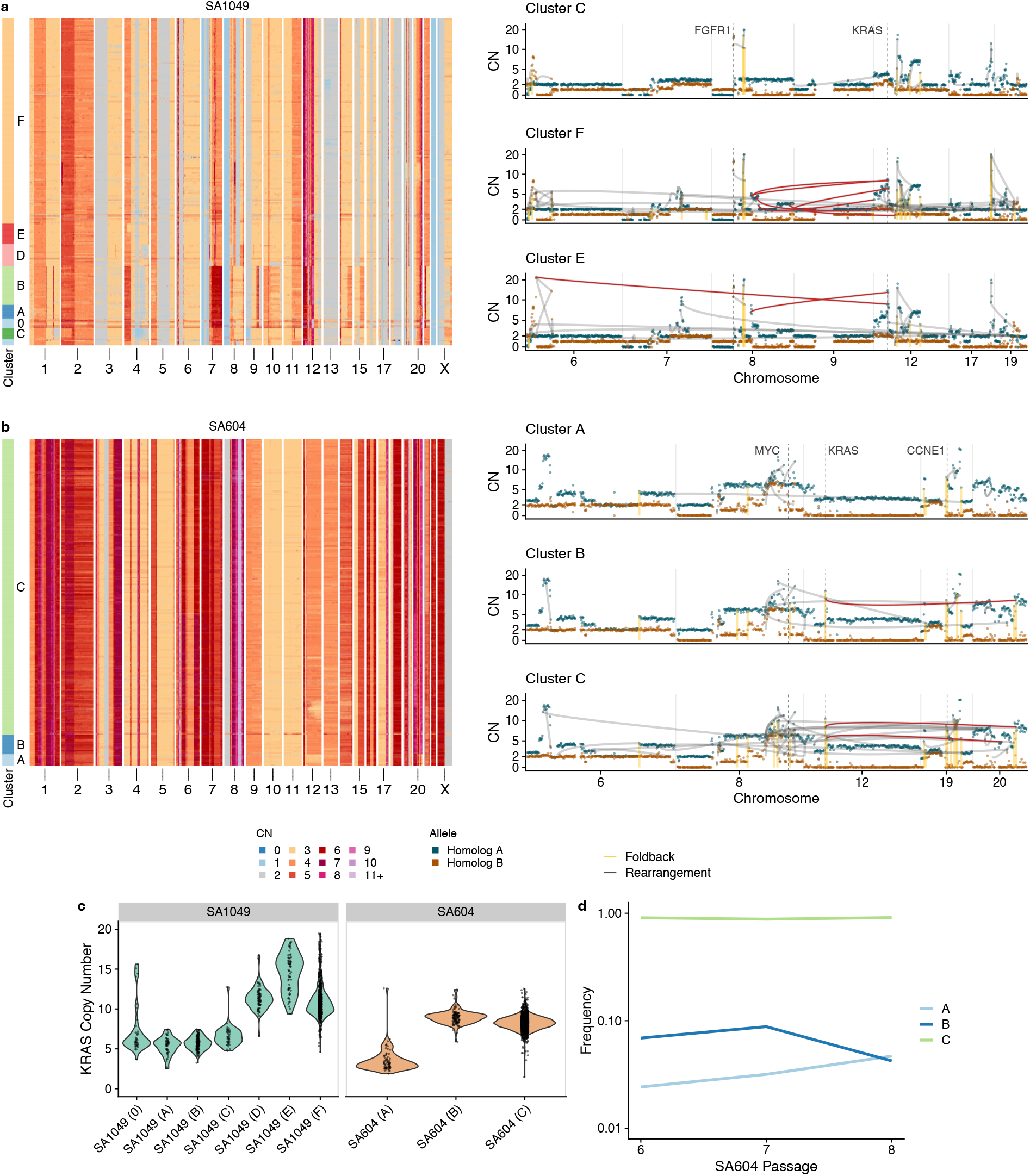
Diversity in complex interchromosomal high level amplifications. a) Left, total copy number heatmap in SA1049 with clusters highlighted with the coloured bar on the left. Right, average allele specific copy number for 3 different clusters overlaid with structural variants. Grey lines indicate links between 2 bins, yellow vertical lines show foldback inversion breakpoints b) same as a) for SA604. Red lines highlight subclonal interchromosomal rearrangement amplifications of interest. c) KRAS copy number per cluster in SA604 and SA1049. d) Frequency of clusters across passage number in SA604.

## Discussion

In this study we reveal substantial genomic diversity in high grade serous ovarian cancers, breast cancers and genetically engineered mammary epithelial cells at haplotype specific resolution. We find evidence of ongoing instability that is distributed uniformly across the genome and were able to estimate rates of chromosomal gains, losses and LOH. Genomic regions that were identified to be at high frequency recurrently across samples are likely to be under positive selection and contribute to tumour progression. Notably, these included focal amplifications of oncogenes such as KRAS, CCNE1 and MYC. Meanwhile, we suggest that pervasive parallel copy number events^10,14,29^ in general are a consequence of underlying levels of instability rather than positive selection. On the other hand, how rarer events modify tumour cell fitness remains uncertain, and will require integration of single cell data with evolutionary models of genomic instability ^30,31^.

Integration of haplotype specific copy number with rearrangement breakpoints allowed us to gain mechanistic insight into the processes generating ITH in single cells. We demonstrate that breakage fusion bridge cycles are a frequent source of genomic diversity and can explain variability in oncogene copy number between cells. Serially passaged PDX models were consistent with BFBC initiation followed by chromosomes undergoing progressive diversification over a few cell divisions until chromosome ends stabilized^17^. Strikingly, we also observed that complex interchromosomal high-level amplifications were also variable between subclones. Complex interchromosomal events are thought to derive from catastrophic genome shattering events^32^. Our time series data point to the possibility that multiple similar but distinct subclones are generated following such an event, as cells attempt to repair their genomes over consecutive cell divisions. In general, subclones containing high level oncogene amplifications had higher clonal frequencies relative to wild-type or low level amplifications (when present), consistent with these amplifications providing a fitness advantage to cells. How subtle differences in amplitude may alter fitness remains unclear however. We suggest that co-existing clones with varying levels of oncogenic amplifications could be exploited as early warnings of phenotypic transformation to a more aggressive state. Our data support the notion that recurrently amplified regions of the genome in breast and ovarian cancers such as at the *PIK3CA*, *CCNE1* and *KRAS* loci have their etiologic origins in BFBC-like processes. Longer read sequencing of clonal haplotypes coupled with genome graph analysis will help to further resolve the mechanistic underpinnings of these events.

Two recent haplotype specific inference methods have been developed for use with the 10X CNV assay^14,15^. These methods also use either haplotype block counts or SNPs genotyped in single cells, differently to these methods, `schnapps` uses a hidden Markov model for inference and facilitates integration with single cell RNA sequencing data. `schnapps` also provides an order of magnitude greater resolution than previous methods (0.5Mb vs 5Mb), enabling reconstruction of the evolution of focal high level amplifications, complemented by integration with subclonal structural variants. `schnapps` also serves as a general toolkit to analyse single cell genomes and includes functionality for clustering, plotting and multimodal integration with scRNAseq. We foresee this toolkit to be a valuable community resource as single cell whole genome sequencing becomes more widely available.

In summary, our study shows how haplotype resolved copy number at single cell resolution can be used to infer instability rates, dissect complex structural rearrangements and identify parallel copy number events. As cohorts of patients profiled at single cell resolution become larger and high throughput methods are applied throughout different stages of disease progression^33^ and across space and time^34^, we envisage that these approaches will enable accurate tracking of the evolutionary history of cancer haplotypes and high resolution characterization of intra-tumour heterogeneity across genomically unstable tumours.

## Methods

### Allele and haplotype-specific copy number in scDNAseq

Previously, we reported allele specific copy number at the level of clones, groups of cells with similar total copy number profiles^11^. This was done by aggregating haplotype block counts within clusters and applying a hidden markov model to infer the most probable state. In this study we extended this approach to the single cell level and also introduce the ability to identify “haplotype specific copy number”. With “haplotype specific copy number number” we can identify cells with the same total copy number but with different allelic combinations. We also leverage haplotype specific copy number to trace the history of complex genomic rearrangements.

First, we’ll summarize the challenges of inferring allele specific copy number in single cells and provide a descriptive overview of our approach. The majority of copy number analysis in single cells works by leveraging differences in read depth across the genome. This is particularly important in sparse single cell approaches such as DLP+ where coverage is of the order 0.01-0.1X. Inference of allele-specific copy number however requires an additional measure of allelic imbalance, in bulk sequencing, this is typically inferred from read count ratios of heterozygous SNPs. This information is very sparse in low coverage single cells, to boost the signal we can infer haplotype blocks from a paired normal sample and then genotype the blocks in single cells. With this in mind, rather than estimating allele-specific copy number using read depth and haplotype counts jointly we decided to leverage the read counts to compute total copy number as we have done previously^11^ and then use allelic imbalance to estimate the allele-specific copy number post hoc. In essence, we assume the total copy number is correct and use this as an input into our allele-specific copy number inference. Validation of our inferences using the BAF distribution per state, matched bulk whole genome sequencing and somatic SNVs present in single cells, confirms that this approach produces reliable estimates.

In this section we’ll describe in detail the schnapps algorithm. The input to our algorithm is total copy number estimates in bins across the genome and haplotype counts per cell. We define the allele-specific state as follows: A|B where A and B are the copy number of the two alleles. The total copy number T is given by A + B, therefore both A and B <= T. Inferring the allele-specific state amounts to identifying the copy state of one of the two alleles. We define the “B allele frequency” as B / (B+A). In most cases, B will be the minor allele across the whole tumour population but our approach does not guarantee this. We note that this is different to how this type of analysis is performed and the data is typically presented in bulk tumour genome sequencing where often both B/(A+B) and A/(A+B) are plotted, resulting in the characteristic split BAF plots in regions of allelic imbalance. As will become apparent, analyzing one of these values rather than both makes distinguishing haplotype-specific copy number intuitively easier. We note that we could in principle use mirrored BAF as is often done in bulk whole genome sequencing and define B as the minor allele in all cases. This is a simpler approach, but does not allow for identification of parallel copy number events and phasing alleles into homologous chromosomes.

We first need to phase the alleles identified in the haplotype blocks into one of the two “tumour” alleles (*A,B*). For the purposes of describing the algorithm, we’ll denote the counts of each “block allele” as (*C_h_*,*D_h_*), and the counts of the phased alleles as (*C_A_*,*D_B_*). For each haplotype block in each cell we get the number of counts assigned to (*C_h_*,*D_h_*) respectively. Our challenge is to identify for each haplotype block how (*C_h_*,*D_h_*) relates to (*A*,*B*), that is we wish to know the phase *P_i_* of each haplotype block, *i*. This gives the counts of the phased alleles, (*C_A_*,*D_B_*). To do this we note that cells will share copy number events and thus we can leverage information across cells to identify block alleles that shift in frequency together. For example, a chromosome undergoing loss of heterozygosity will completely lose either the maternal and paternal allele, thus any block alleles within the LOH event that contain non zero counts must necessarily be phased together. As a first approximation we first assign the B allele to be the minor allele across all cells:

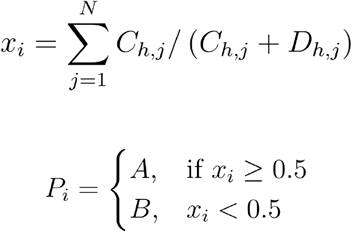

When a particular region of the genome is in a balanced state across all cells, distinguishing A and B is not possible. In this case, (*A*,*B*) will be assigned randomly due to stochastic fluctuations in read counts.

After this initial phasing assignment we then merge the phased haplotype block counts within bins and compute a BAF value for each bin in each cell:

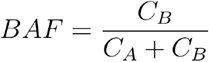

With these values we then used a HMM to compute the optimal allele specific state. We used a beta-binomial emission model and the Viterbi algorithm to compute the optimal B-allele state. Given observed total copy number, *T* unobserved B-allele copy number *B*, B-allele counts and total counts *C_T_* the likelihood is given by

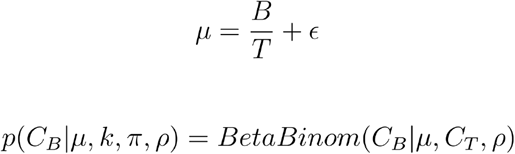

Where *ϵ*is an error term included to account for noise in the data, which we set to 0.01 in the first instance. This is particularly important in LOH states, where for example due to noise the BAF is rarely exactly 0.0. *ρ*is the degree of overdispersion in read counts, which can be inferred from the data, when *ρ* → 0.0 we recover a Binomial likelihood.

We used the following transition matrix setting *δ* = 0.95, favouring self-transitions.

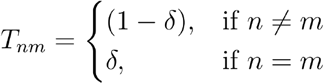

Following the above steps gives us our first allele-specific assignment. However, this assignment can have some issues due to inaccurate phasing from the first phasing step. Because our initial phasing uses the minor allele across all cells, if there are a number of overlapping events in different cells at different proportions we sometimes find implausible results, where for example a cell will switch phase in the middle of a chromosome, see **Supplementary Figure 11** for a diagram showing how this can arise. To avoid this, we go through a second round of phasing and inference. We assume that the most accurate phasing should favour results that minimize the number of apparent switches in phasing. To do this, for each chromosome we cluster BAF values from step 1, and then identify the cluster with the largest amount of imbalance in each chromosome. Using this cluster as an anchor we then define the B allele as the minor allele of cells within this cluster. Clustering is performed using umap and hdbscan as described below. Haplotype blocks are then reassigned their phase relative to this cluster. Following this reassignment, we then rerun the HMM. Prior to running the HMM, we also take advantage of this 2 step process to infer *ϵ* and *ρ*directly from the data and assess statistical support for the Binomial vs BetaBinomial likelihood model. *ϵ* is computed from the average BAF of states assigned as homozygous, we compute Tarones z-score to assess statistical support for BetaBinomial model ^35^. If we find support for a BetaBinomial model (z>5), *ρ* is then computed using maximum likelihood estimation. The HMM is then rerun with these input parameters and new phasing producing the final allele-specific assignment.

When we use allele specific copy number we use mirrored BAF and assign B to always be the minor allele in every cell, such that A >= B. Therefore a state such as 1|2 will become 2|1. We used allele specific copy number in figure 1, but for the remainder of the analysis we used haplotype specific copy number.

`schnapps` is available as an R package at https://shahcompbio.github.io/schnapps/. As well as the allele specific copy number algorithm, schnapps includes a large number of functions for plotting copy number profiles and heatmaps, clustering cells, integrating with scRNAseq (see below) and performing QC. A number of tutorials accompany the package at the above URL describing this functionality.

### Comparison to other methods

Recently, two other methods (CHISEL^14^ and Alleloscope^29^) were published that infer allele specific copy number from sparse single cell sequencing data. These methods were applied to data generated from the 10X CNV assay. As is the case with schnapps, CHISEL uses haplotype block counts for inference, while Alleloscope uses the raw SNP counts. Both methods use an approach based on clustering BAF and read depth jointly to assign allele specific copy number. Differently to these methods, schnapps directly models the read counts of haplotype blocks (with and without overdispersion) using a hidden markov model and uses a clustering approach to phase haplotypes. The resolution of schnapps is 0.5Mb whereas CHISEL uses 5Mb bins. Alleloscope segments the genome before inference, so resolution will be a function of the segmentation. Differently to the other methods, schnapps also provides an approach for integration with single cell RNA seq, a feature unique to `schnapps`. Alleloscope on the other hand is unique in that it provides methods to integrate single cell DNA sequencing with single cell ATAC-seq.

### Experimental methods

Detailed description of the data generation methods are described in Funnell *et. al.*^20^. Including generation of engineered cell lines, xenografting, tissue processing, single cell whole genome sequencing and bulk whole genome sequencing.

### DLP+ whole genome sequencing quantification and analysis

Single cell copy number, SNV, SV and haplotype block calls were generated using our previously described approach^11^, except that BWA-MEM was used to map DLP+ reads to the hg19 reference genome. The genome was segregated into 500 kb bins, and GC-corrected read counts were calculated for each bin. These read counts were then input into HMMCopy to produce integer copy number states for each bin^36^.

To detect SNVs and SVs in each dataset, reads from all cells in a sample were merged to form “pseudobulk” libraries. SNV calling was performed on these libraries individually using MutationSeq^37^ (filtered by probability threshold = 0.9) and Strelka (filtered by score > 20) ^38^. Only SNVs detected by both methods were retained. For each dataset, the union of SNVs was aggregated, then for each cell and each SNV, the sequencing reads of that cell were searched for evidence of that SNV. SV calling was performed in a similar manner, by forming pseudobulk libraries, then running LUMPY^39^ and DESTRUCT^40^ on each pseudobulk library.

To call haplotype blocks we identified SNPs from the 1000 genomes phase 2 reference panel in matched normal sample. An exact binomial test was used to identify heterozygous SNPs which were then input into shapeit to identify haplotype blocks^41^. SNPs used in the haplotype block inference were then genotyped in individual cells producing per cell haplotype block counts that could be used for allele specific copy number inference with schnapps.

### Bulk whole genome sequencing

Bulk whole genome sequencing data was generated from matched primary samples from all patients. Reads were aligned to hg19 using BWA-MEM. Genome wide allele specific copy number was called using Remixt^42^ with default parameters.

### DLP+ data filtering

Cells were retained for further analysis if the cell quality was at least 0.75 ^11^, and they passed both the s-phase and contamination filters. The contamination filter uses FastQ Screen^43^ to tag reads as matching human, mouse, or salmon genomes. If >5% of reads in a cell are tagged as matching the mouse or salmon genomes, then the cell is flagged as contaminated. The s-phase filter uses a Random Forest classifier and removes cells where s-phase is the most probable state ^11^. Samples were also filtered to remove small numbers of contaminating diploid cells. We also used the procedure outlined in to further filter out any replicating cells that were missed by the s-phase filter. As the allele specific copy number inference requires cell level haplotype block counts per cell, we additionally filtered out any cells that contained < 100 haplotype block counts.

### 10X scRNAseq data generation

184hTERT cells were cultured as previously described ^34,44^ in MEBM (Lonza) supplemented with the SingleQuots kit (Lonza), 5 μg/ml transferrin (Sigma-Aldrich) and 10uM isoproterenol (Sigma-Aldrich). Cells were pelleted and gently resuspended in 200ul PBS followed by 800ul 100% methanol and incubation at −20C for 30mins to fix and dehydrate cells. Cells were then pelleted and resuspended in 0.04% BSA/PBS and immediately loaded onto a 10x Genomics Chromium single-cell controller targeting 3,000 cells for recovery. Libraries were prepared according to the 10x Genomics Single Cell 3′ Reagent kit standard protocol. Libraries were then sequenced on an Illumina Nextseq500/550 with 42-bp paired end reads, or a HiSeq2500 v4 with 125-bp paired end reads.

### 10X scRNAseq data analysis

The pipeline is built using 10X Genomics *Martian* language and computational pipeline framework. *CellRanger* software (version 3.1.0) was used to perform read alignment, barcode filtering, and UMI quantification using the 10x GRCh38 transcriptome (version 3.0.0) for FASTQ inputs. *CellRanger* filtered matrices are loaded into individual *Seurat* objects using the *Seurat* R package (version 3.0.1)^45,46^. The resulting gene by cell matrix is normalized and scaled for each sample. Cells retained for analysis had a minimum of 500 expressed genes and 1000 UMI counts and less than 25% mitochondrial gene expression. Cell cycle phase was assigned using the *Seurat*^46^ CellCycleScoring function. *Scrublet*^47^ (version 0.2.1) was used to calculate and filter cells with a doublet score greater than 0.25.

### Allelic imbalance in scRNAseq

We called heterozygous SNPs in the scRNAseq data using cellSNP^48^. As input, we used the same set of heterozygous SNPs identified in the scDNAseq and corresponding normal sample for each sample. The liftover script provided in cellSNP was used to lift over SNPs from hg19 to hg38. Following genotyping, we phase the SNPs using the phasing information computed from the allele specific inference in the scDNAseq. As SNP counts are much more sparse in scRNAseq vs scDNAseq (∼2 orders of magnitude lower), we aggregated counts across chromosome arms, computing the BAF for each arm. We then generated a cell by chromosome arm BAF matrix and incorporated this into our gene expression Seurat objects. Functionality to map scDNAseq to scRNAseq and call allelic imbalance are provided in `schnapps`. Density of the cells with loss of different haplotypes were plotted using the Nebulosa R package^49^.

### Phylogenetic analysis

We used a previously described phylogenetic method sitka to generate single cell trees for each sample^19^. Sitka uses breakpoints (also referred to as changepoints) in copy number across the genome as phylogenetic characters to construct the evolutionary relationships. Rather than use total copy number as previously described, here we used haplotype specific copy number. To do this, we enumerated breakpoints on each haplotype. For example a loss of haplotype A will have a separate breakpoint feature than a loss of haplotype B even if the genomic position of the losses are identical. This allows for phylogenetically distinguishing parallel evolutionary events. There can be some cell-to-cell variability in breakpoints that is technical rather than biological, due to for example fluctuations in read and SNP counts. To mitigate the influence of this variability, we averaged the copy number profiles in 3Mb windows, ensuring consistent breakpoints across cells as much as possible. `sitka` was run for 3,000 chains and a consensus tree was computed for downstream analysis.

### Clustering copy number profiles

To cluster copy number profiles we used UMAP dimensionality reduction followed by HDBSCAN ^11,50,51^. This is implemented within schnapps (function `umap_clustering`) with following default parameters:

- Distance metric: correlation
- Number of neighbours: 10
- Minimum distance = 0.1
- Minimum number of points in cluster: 30

### Pseudobulk allele specific copy number profiles

In numerous places in the text we construct “pseudobulk” allele specific copy number profiles either across all cells in a sample or subsets of cells that share some features of interest. To do this we group the cells of interest and then compute an average profile by taking the median values of copy number and BAF and the mode of the allele specific state. The function `consensuscopynumber` provided in `schnapps` was used for this.

### Chromosomal event rates and LOH rate analysis

To compute chromosomal event rates we enumerated the number of events from our single cell phylogenies using parsimony based ancestral state reconstruction. We first computed whole chromosome level consensus copy number profiles for each cell, allowing us to assign chromosome level states to each tip (cell) in the phylogeny. We defined states relative to cell ploidy, identifying for each arm whether the chromosome was gained or lost and whether the chromosome was homozygous. For each chromosome, cells can have one of two possible states for each class of interest: (Gain, not gained), (Loss, not lost), (LOH, not LOH). By casting the problem as reconstructing the ancestral states within the phylogeny we can then compute the number of transitions between these states that most parsimoniously explains the phylogenetic tree. We used a simple transition matrix where transitions between states incurs a cost of 1. Ancestral state reconstruction then amounts to finding the reconstruction that minimizes this cost, we refer to this cost as the parsimony score. The event frequency per sample per chromosome is then calculated by dividing the parsimony score (number of events) by the number of cells. We used castor in R to perform the ancestral state reconstruction ^52^. As we were interested in LOH events that were not just due to losses resulting in a single copy, we removed LOH events where the state was 1|0 from this calculation. The units of this quantity is the number of events per chromosome per cell division assuming no cell death. It’s possible (perhaps likely) that many cells get chromosomal gains or losses but then die, we of course never sample such cells and our phylogenetic tree reconstructs ancestral relationships between cells that survive and that we sample. This value is therefore likely to be an overestimation of the true cell division rate if there is considerable cell death. It is challenging to decouple the death rate of cells from the true event rate per cell division, see Werner *et. al.* for a similar problem^53^. To get a summary value for each sample we took the mean of the chromosome level estimates per sample, this value is what is used in Figures 2 and 3.

### Identification of parallel copy number events

Parallel copy number events were defined as genomic regions greater than 4Mb where gain or loss of both the maternal and paternal haplotype was observed in more than 1% of cells. This calculation will be influenced by the number of cells sequenced so in order to compare the number of parallel events across tumours we divided this number by the number of cells.

### Identification of BFBC

BFBC have a number of characteristic features which we attempted to identify in our single cell data: i) staircase copy number patterns ii) foldback inversion rearrangements coincident with copy number changes and iii) amplifications adjacent to losses. Amplifications may also appear at the terminal end of a chromosome when telomeres are short. **Supplementary Table 2** summarizes the evidence for BFBC in terms of these features for each example described in the main text or included in the supplementary figures. When an amplification is adjacent to a loss, BFBC would predict that both the amplification and the loss occurs on the same homologous chromosome. Haplotype specific copy number allows for this inference to be made, however in some cases, this information may be ambiguous. In some cases the default output from `schnapps` may assign the gain and the loss to different haplotypes, this is because in the absence of a cluster of cells with different copy number schnapps will assign the “B” allele to be the minor allele in the whole tumour population. In these cases we first looked for rare cells that had whole chromosome losses which would provide unambiguous phasing information (we assume the whole chromosome loss was a single event and affected the same homolog). In many cases we could identify such cells and adjust the phasing accordingly. This was the case for both SA535 and SA1035, the 2 examples we looked into in detail in Figure 5. To group cells into clusters we used the UMAP + HDBSCAN clustering approach outlined above but only clustered using bins within the chromosome of interest. Clustering is therefore chromosome specific. For each cluster we constructed consensus haplotype specific copy number profiles and assigned rearrangement breakpoints to clusters if any cell within the cluster had evidence of the breakpoint.

### Identification of interchromosomal high level amplifications

We used the rearrangement breakpoints to identify samples where high level amplifications were linked across chromosomes. We clustered cells only including bins that were part of the chromosomes of interest. In sample SA1049, chromosomes 6, 7, 8, 9, 12, 17 and 19 were used for clustering. In SA604 we were particularly interested in the co-amplification of chr12 and chr20 so restricted the clustering to those chromosomes only.

### PCAWG data

Copy number calls from PCAWG were downloaded from the ICGC portal (https://dcc.icgc.org/releases/PCAWG/). We transformed the segmentations into 0.5Mb bins across the genome to facilitate comparison with our single cell data. We filtered the PCAWG data for ovarian and breast cancer types for downstream analysis.

### Recurrent event analysis

To identify recurrent events across the cohort we first classified each genomic bin in each cell into gains, losses and LOH. LOH states include any event that has lost one of the alleles, for example monosomies (ie 1|0), copy neutral LOH (ie 2|0) and regions that are also gained (ie 3|0) were all included under LOH. Therefore some bins will be classified as both LOH and loss or LOH and gain. Bins were assigned to be gained or lost relative to cell ploidy. After assigning these states we then computed the cancer cell fraction, *f_t_* of each type in bins across the genome:

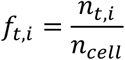

Where *n_t,i_* is the number of cells with event type *t* in bin *i*, and *n_cell_* is the total number of cells in the sample. To look at recurrency across samples we then took these values and computed the fraction *F*of samples that had an event in bin *i* with *f_t,i_* greater than some cutoff *X*.

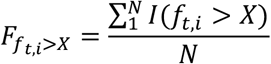

Where *N* is the number of samples and *I* is the indicator function. In Figure 2 we used cutoffs, *X* of 0.01 and 0.95.

To investigate how the prominence of focal alterations around oncogenes changes as a function of CCF we calculated the ratio, *R_g_* between *F* around the locus of interest to the average across the whole chromosome:

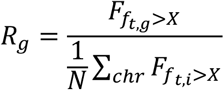

Where *N* is the number of bins in the chromosome of interest and *g* is a gene of interest. We calculated *R_g_* in 250 oncogenes from the cancer gene census across a range of CCF’s. These were then plotted in **Figure 2d**.

### Statistical tests

To compare the proportion of cells with loss of haplotype A vs B in gene expression clusters we used a proportions test (using `prop.test` in R). Linear regressions use the `lm` function in R. When boxplots are presented in the figures, hinges represent the 25% and 75% quantiles, whiskers are +/− 1.5X inter quartile range.

### Code availability

- Schnapps R package: https://shahcompbio.github.io/schnapps/.
- Analysis scripts and figure generation: https://github.com/marcjwilliams1/schnapps_paper
- DLP+ single cell whole genome sequencing pipeline is available at https://github.com/shahcompbio/single_cell_pipeline
- Whole genome sequencing pipeline: https://github.com/shahcompbio/wgs
- scRNAseq pipeline: https://github.com/nceglia/scrna-pipeline

### Data availability

10X scRNA sequencing data from SA906 is available from the European Genome-Phenome archive (EGAS00001004448). All other data will be made available for controlled access at EGA upon publication.

## Acknowledgments

This project was generously supported by the BC Cancer Foundation at BC Cancer and Cycle for Survival supporting Memorial Sloan Kettering Cancer Center. SPS holds the Nicholls Biondi Chair in Computational Oncology and is a Susan G. Komen Scholar (#GC233085). SA holds the Nan and Lorraine Robertson Chair in Breast Cancer and is a Canada Research Chair in Molecular Oncology (950-230610). Additional funding provided by the Terry Fox Research Institute Grant 1082, Canadian Cancer Society Research Institute Impact program Grant 705617, CIHR Grant FDN-148429, Breast Cancer Research Foundation award (BCRF-18-180, BCRF-19-180 and BCRF-20-180), MSK Cancer Center Support Grant/Core Grant (P30 CA008748), National Institutes of Health Grant’s (1RM1 HG011014-01 & P50 CA247749-01), CCSRI Grant (#705636), the Cancer Research UK Grand Challenge Program, Canada Foundation for Innovation (40044) to SA and SPS. MJW is supported by an National Cancer Institute pathway to independence award (K99CA256508).

## Competing interests

SPS and SA are shareholders and consultants of Canexia Health Inc.

## Contributions

SPS, SA and MJW: project conception, manuscript writing, senior responsible authors; SPS and SA: project supervision and oversight; NR: manuscript writing and editing; COF, FK, HL, TM, PE, DY, BW, JB, JB, JT: tissue procurement, biological substrates, knockout cell line generation and validation, data generation. COF, JB, BW, JB: single cell sequencing; TF, MJW, SS, IVG, AM, ACW, NC, FU: computational biology, data analysis; DL, SB, JP, DG, DA, AM, SL, EH, VB: data processing, visualization;. All authors read and approved the final manuscript.

## Supplementary Figures

**Supplementary Figure 1.**
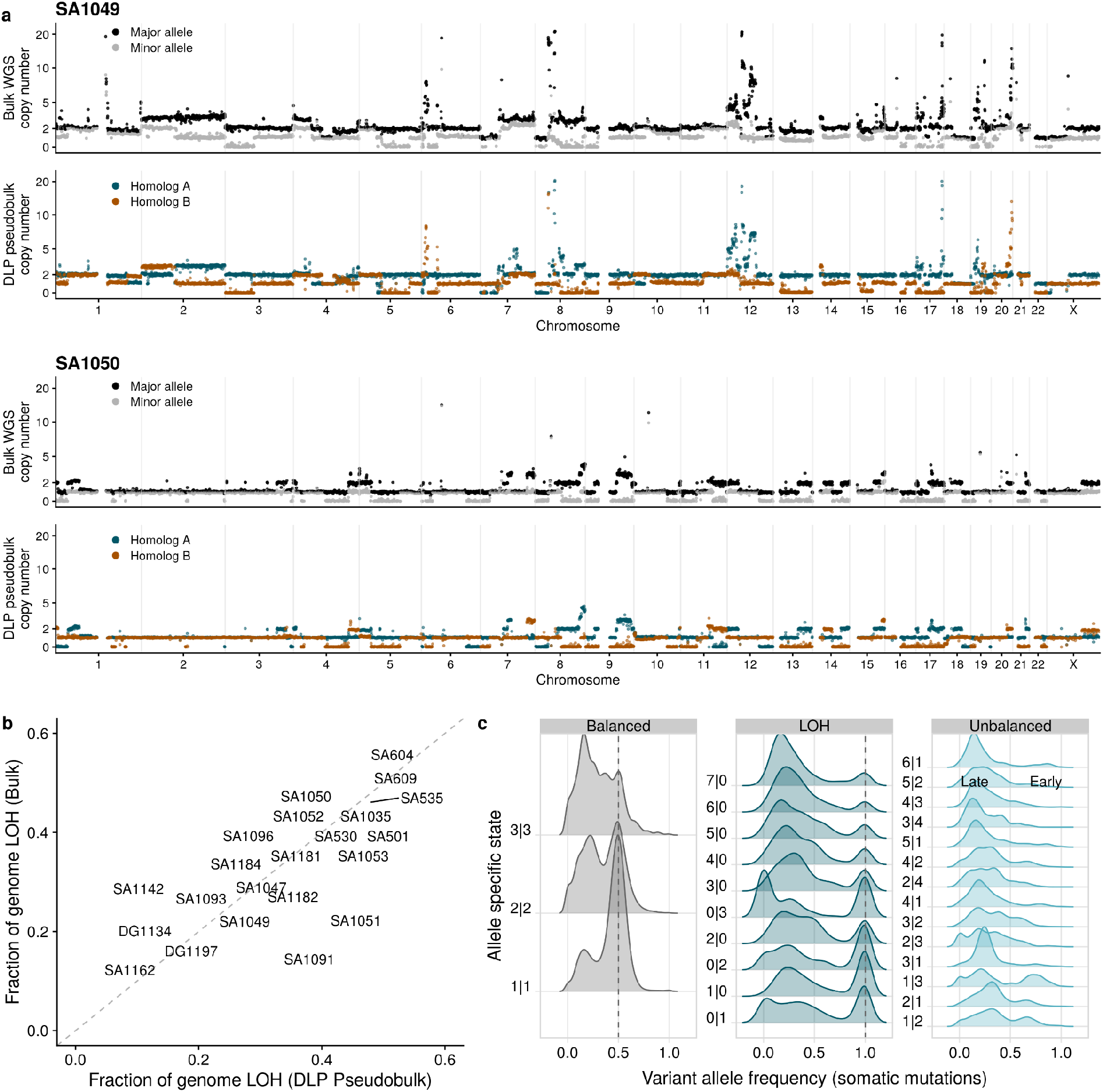
Comparison of bulk whole genome sequencing and DLP. a) Allele specific copy number inferred from bulk WGS using RemixT (top), pseudobulk allele haplotype copy number (bottom) for samples SA1049 and SA1050. b) Fraction of genome inferred to be LOH in pseudobulk DLP vs bulk WGS. c) Density of variant allele frequency of somatic SNVs stratified by allele specific state across all samples.

**Supplementary Figure 2.**
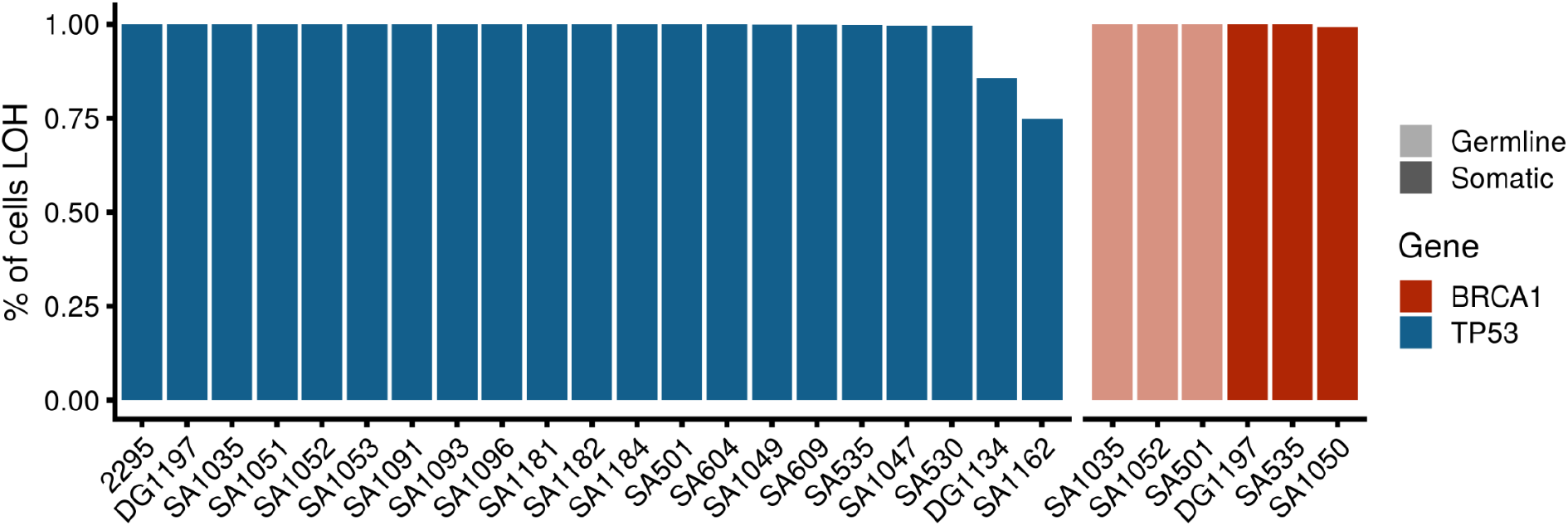
Percentage of cells that are homozygous around the BRCA1 and TP53 locus where we could identify a loss of function mutation. x- axis is the sample ID and y-axis is the % of cells that are homozygous.

**Supplementary Figure 3.**
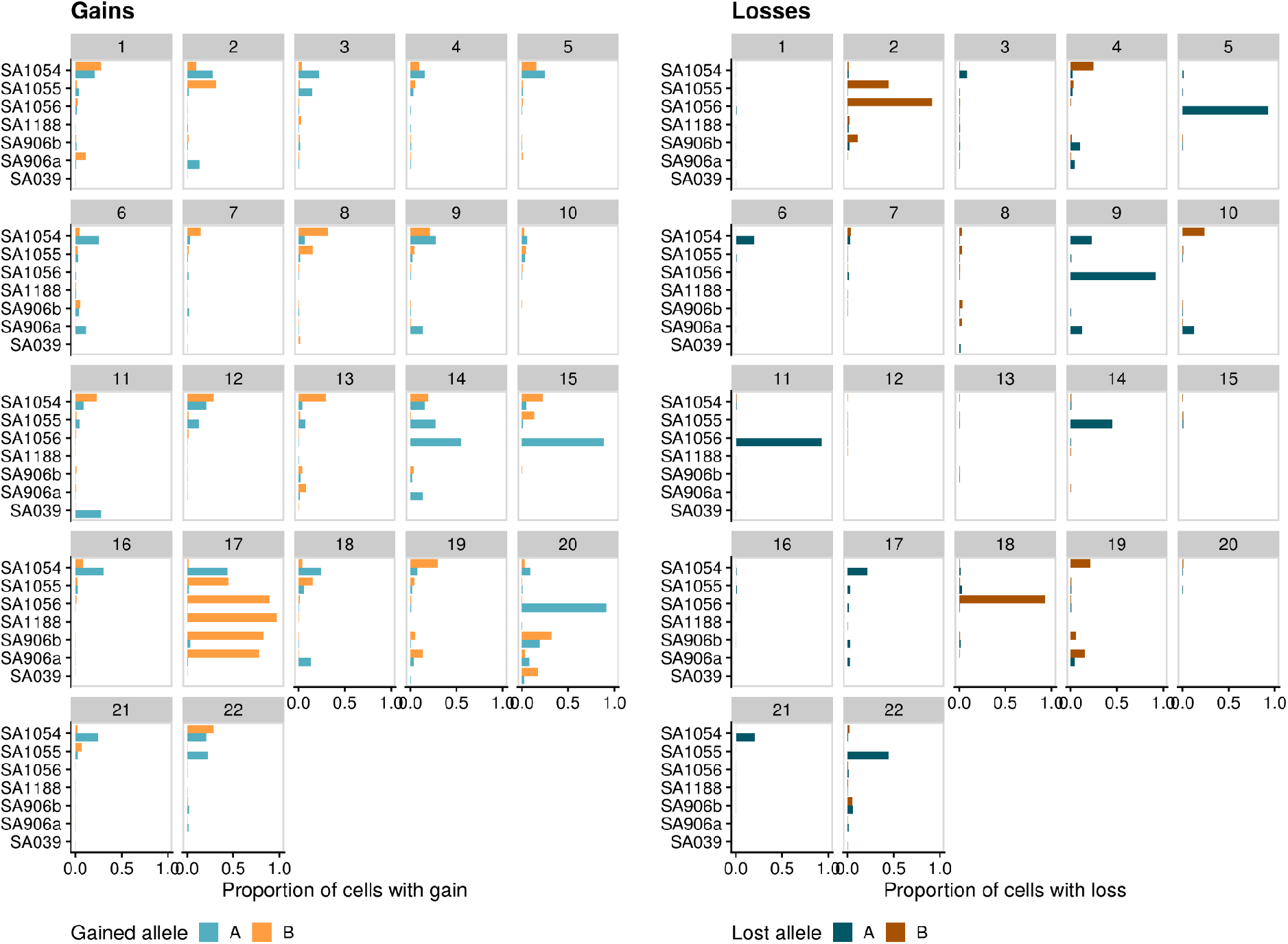
Proportion of cells with gains (left panel) and losses (right panel) of allele A and allele B across chromosomes for each of the engineered cell lines. As these cells share a common ancestor, haplotypes can be phased jointly across all cells so allele A and B are consistent across the different lines.

**Supplementary Figure 4.**
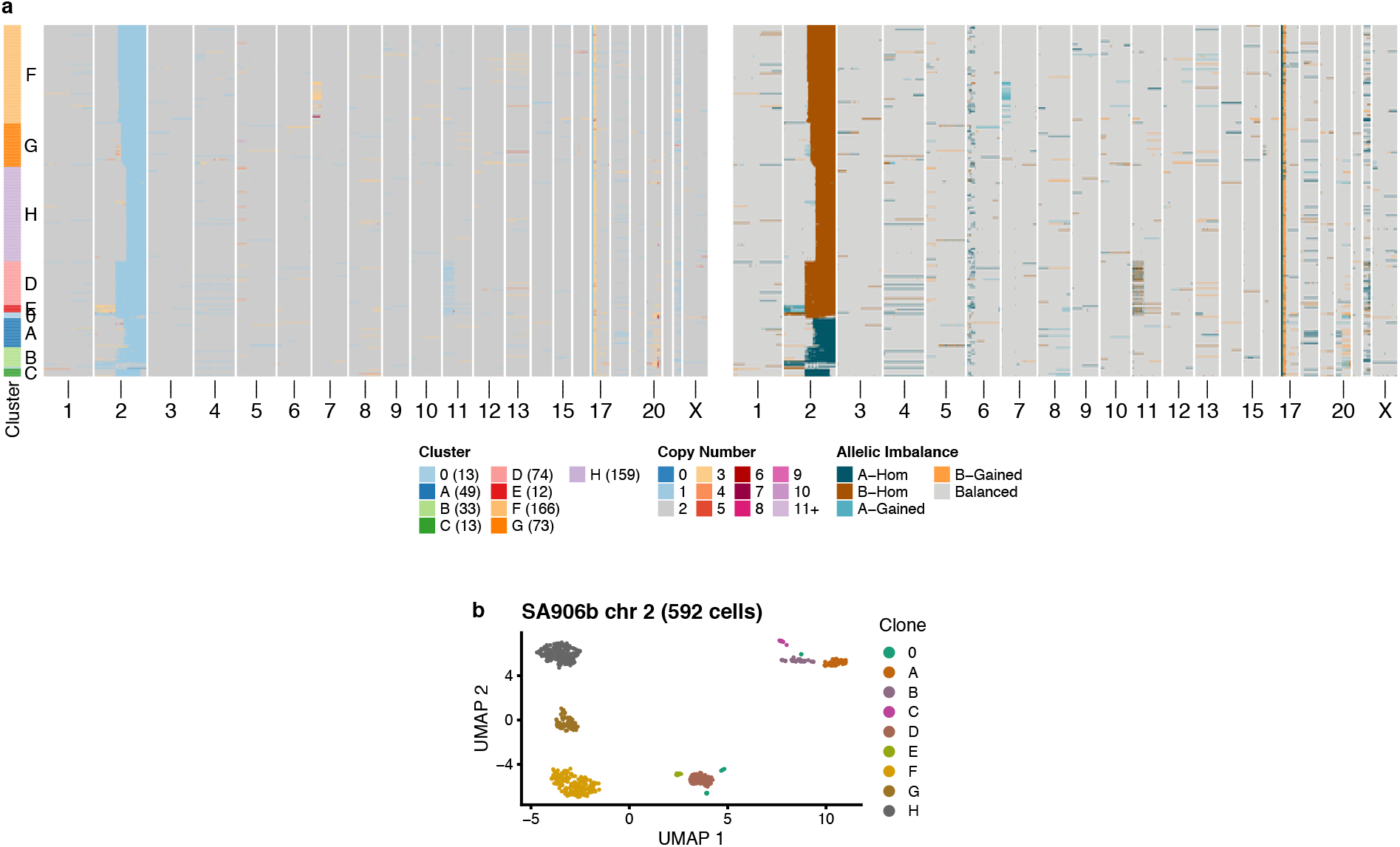
a) Total copy number (left) and haplotype specific copy number of cells in SA906b that have a loss on chr 2q. Each row is a cell and x-axis is genome position. Left track shows groupings into clusters using UMAP and HDBSCAN. Same set of cells shown in Figure 3 is shown here. b) Output of UMAP showing distinct clusters. Points are coloured according to clusters/clones. Same clustering is used her as in Figure 3.

**Supplementary Figure 5.**
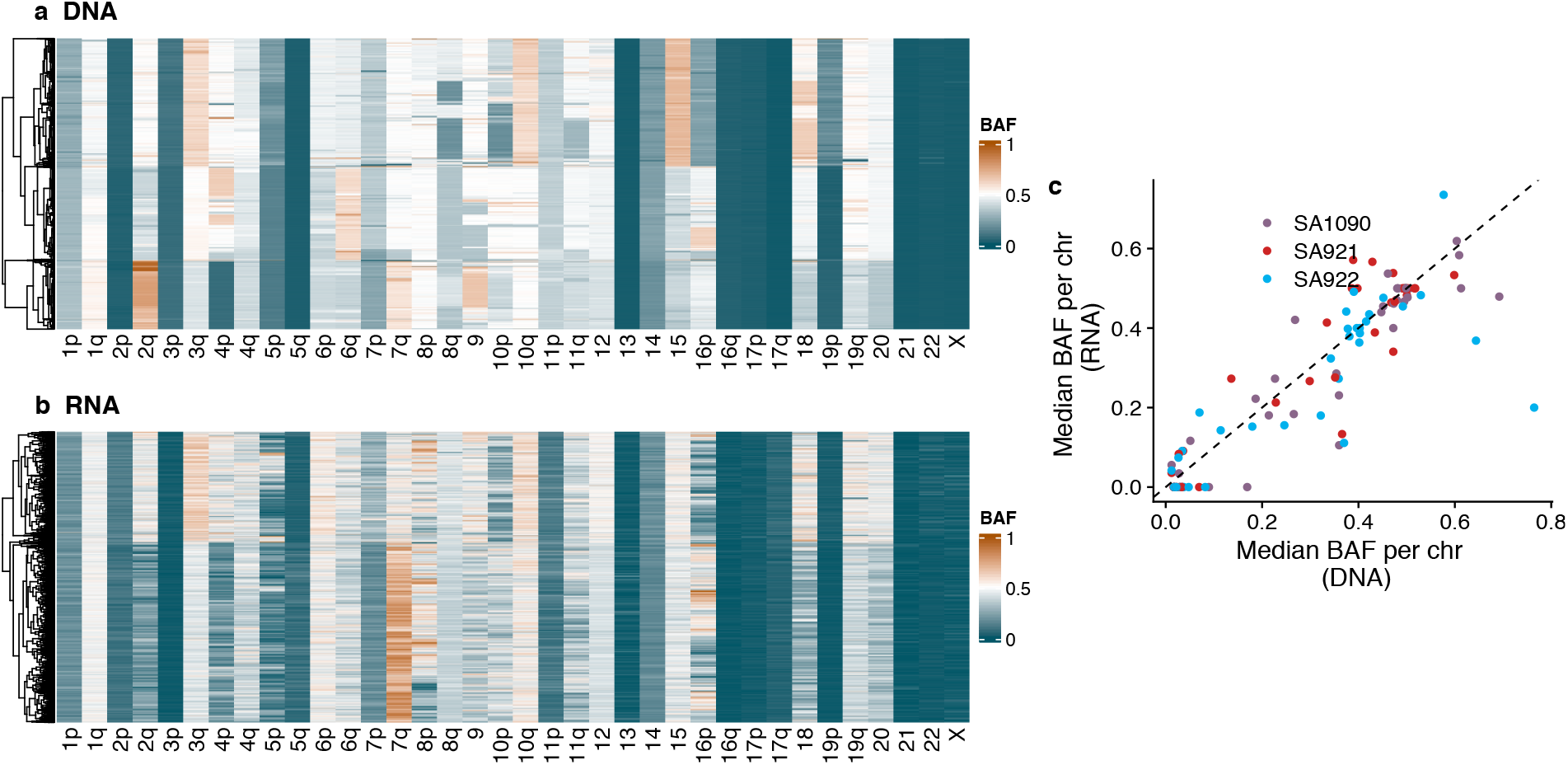
Comparison of allele imbalance per chromosome inferred from single cell DNA a) versus single cell RNA b) in sample cell lines 2295 (same sample used in Figure 1). In a) and b) each column is a chromosome arm and each row is a cell, colours indicate allele imbalance. c) Median BAF per chromosome arm inferred from scDNA vs scRNA, colours indicate the different sites and dashed black line the y=x line.

**Supplementary Figure 6.**
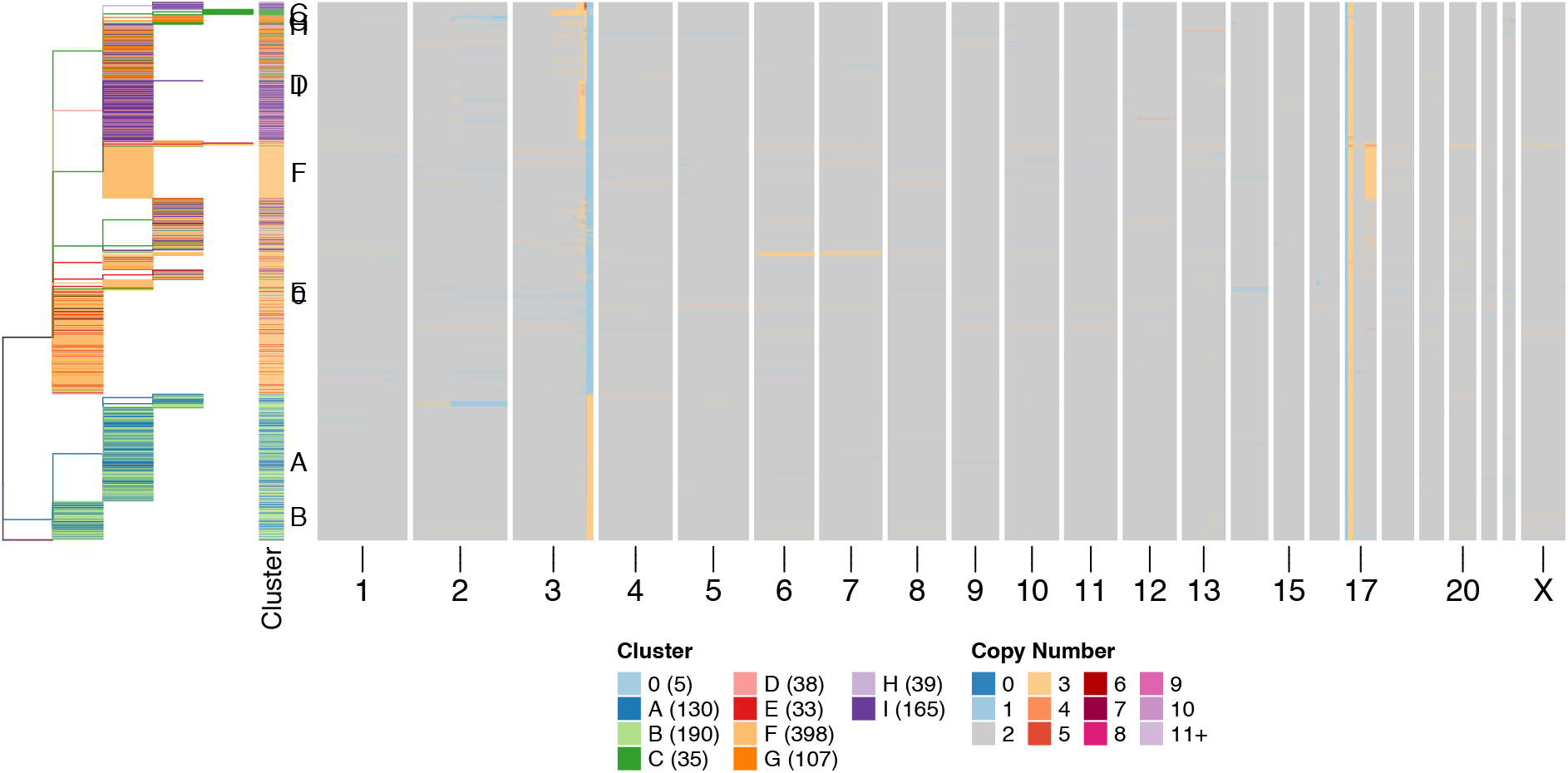
Heatmap ordered by phylogenetic tree for SA1188. Rows are cells and x-axis represent genome position. Tree and cluster track are coloured according to clustering presented in Figure 4, showing that clusters largely group together on the phylogenetic tree and that a split close to the root is present which distinguishes two clades, one clode with an amplification at the end of chromosome 3 and one with a deletion.

**Supplementary Figure 7.**
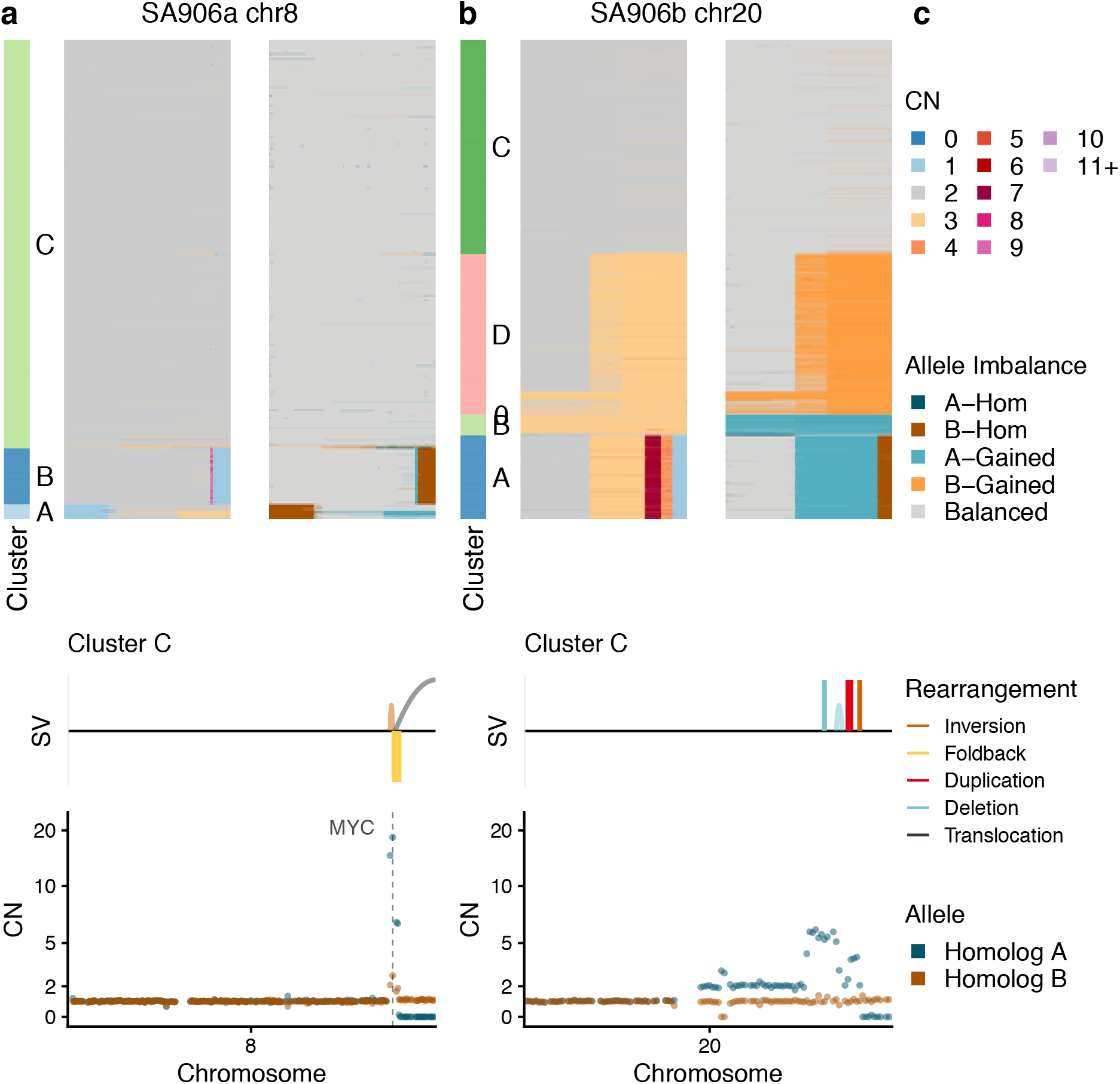
BFBC in htert engineered cell lines. For each panel we show the total copy number on the left, allele specific copy number on the right and zoomed in haplotype specific copy number plot for the cluster containing the BFBC event at the bottom. We show: a) MYC amplification in sa906a and b) chr20 amplification in SA906b

**Supplementary Figure 8.**
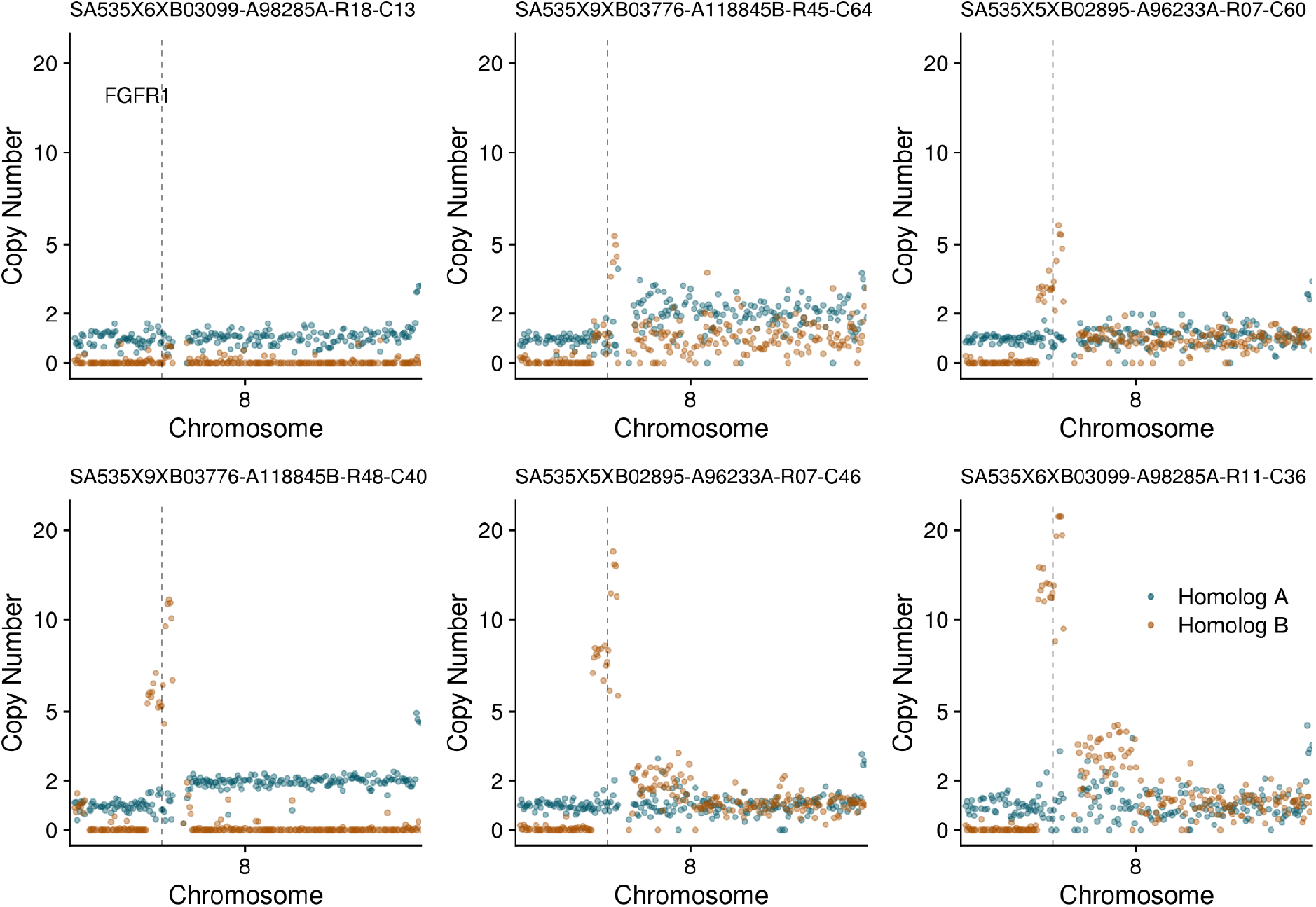
Examples of single cells with progressive amplifications on chr 8p in SA535. Each panel is the haplotype specific copy numbers in chr8 in individual cells. The cell id is given at the top of each panel, the location of FGFR1 is shown with a dashed line.

**Supplementary Figure 9.**
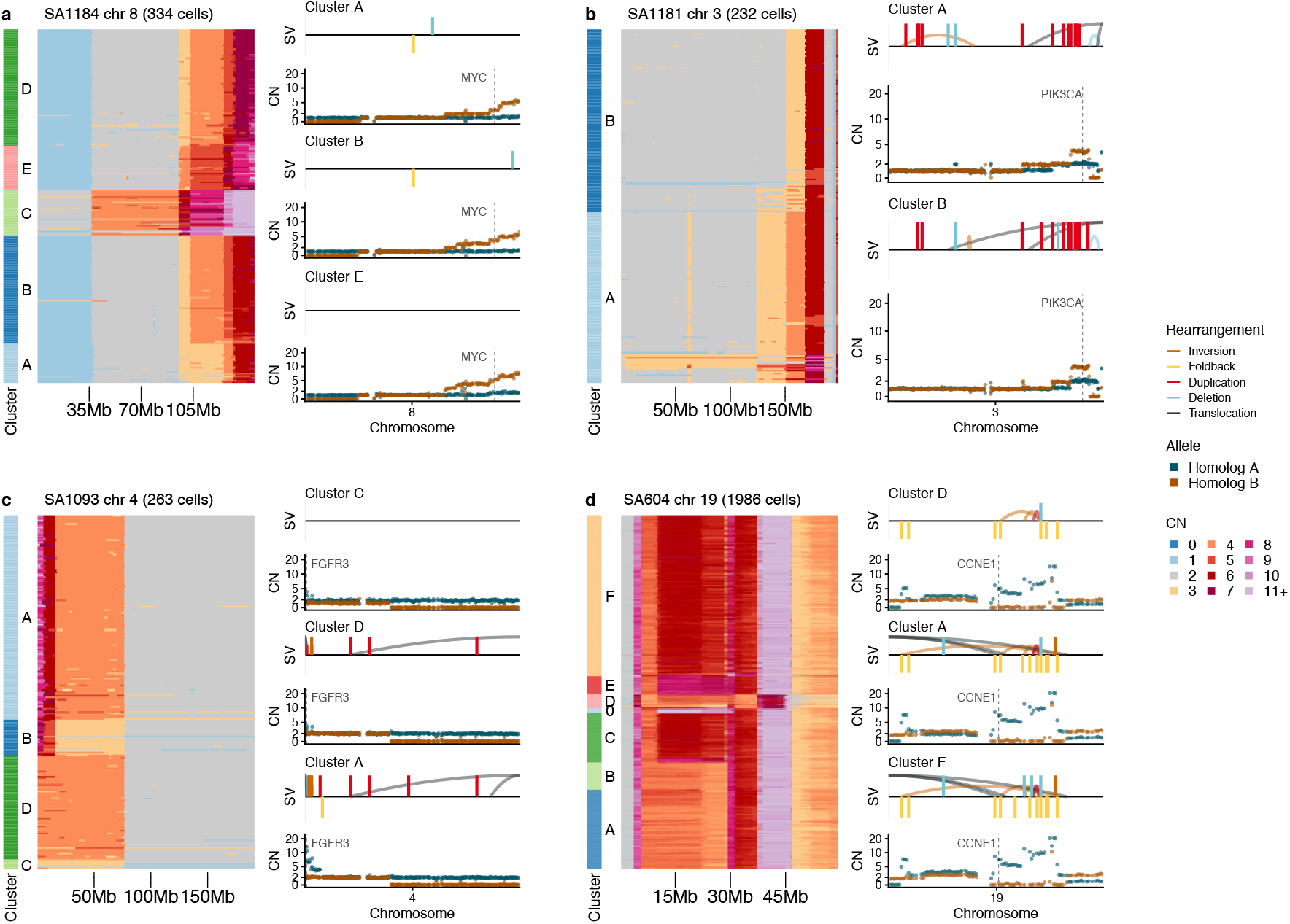
Additional examples of putative BFBC resulting in subclonal and/or variable oncogene amplifications in tumours For each panel we show the total copy number heatmap grouped into clusters with each row being a cell on the left and pseudobulk haplotype specific copy number plots with structural variants from some clusters on the right. The examples shown here are:. a) MYC in SA1184 b) PIK3CA in SA1181 c) FGFR1 in SA1093 and d) KRAS in SA604.

**Supplementary Figure 10.**
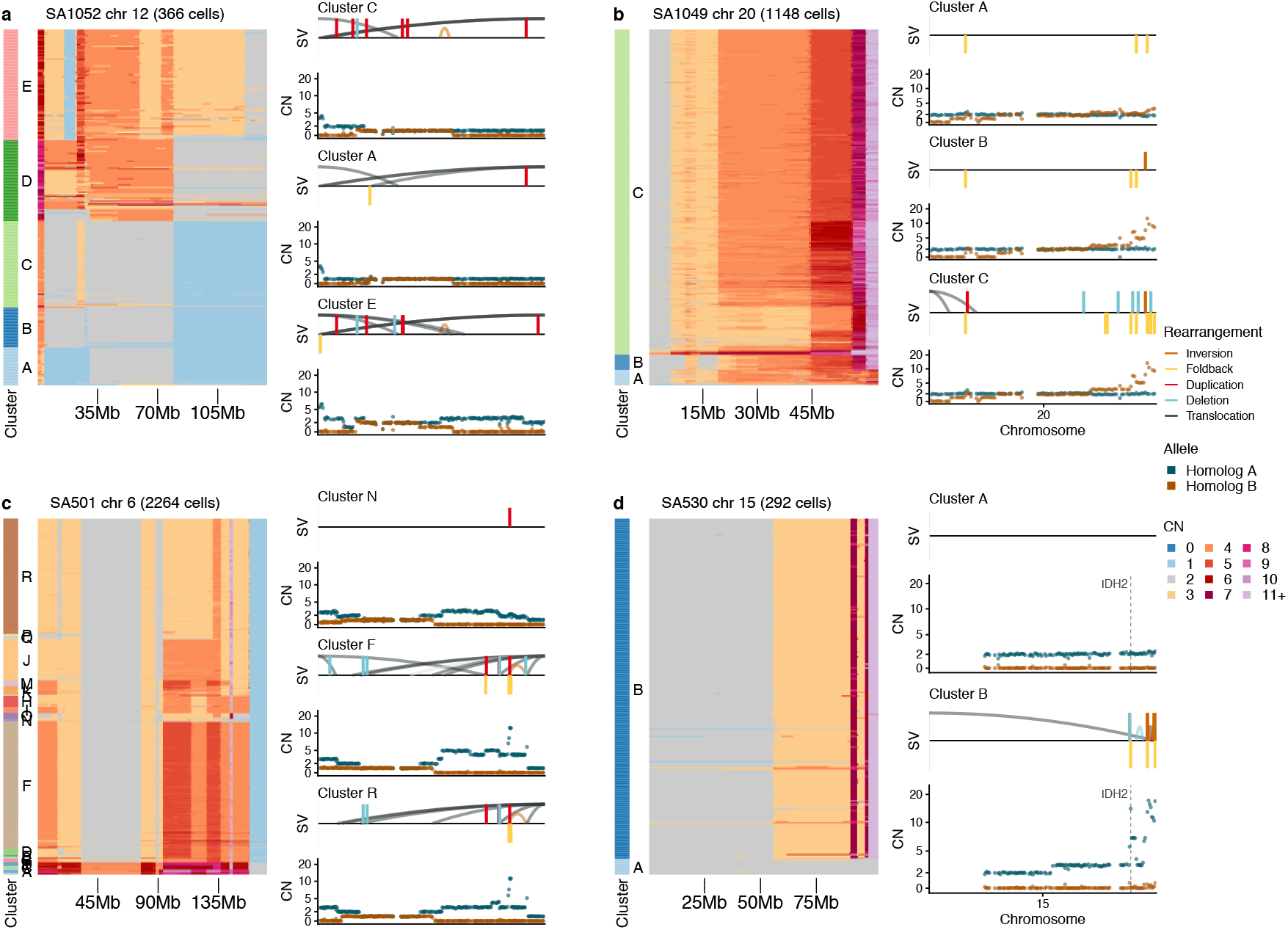
Additional examples of putative BFBC with no associated oncogene amplified. For each panel we show the total copy number heatmap grouped into clusters with each row being a cell on the left and pseudobulk haplotype specific copy number plots with structural variants from some clusters on the right. Examples shown are: a) chr12 SA1052 b) chr 20 SA1049 c) chr 6 in SA501 and d) chr 15 in SA530 chr6. SA530 amplified IDH2 but the significance of this alteration is uncertain.

**Supplementary Figure 11.**
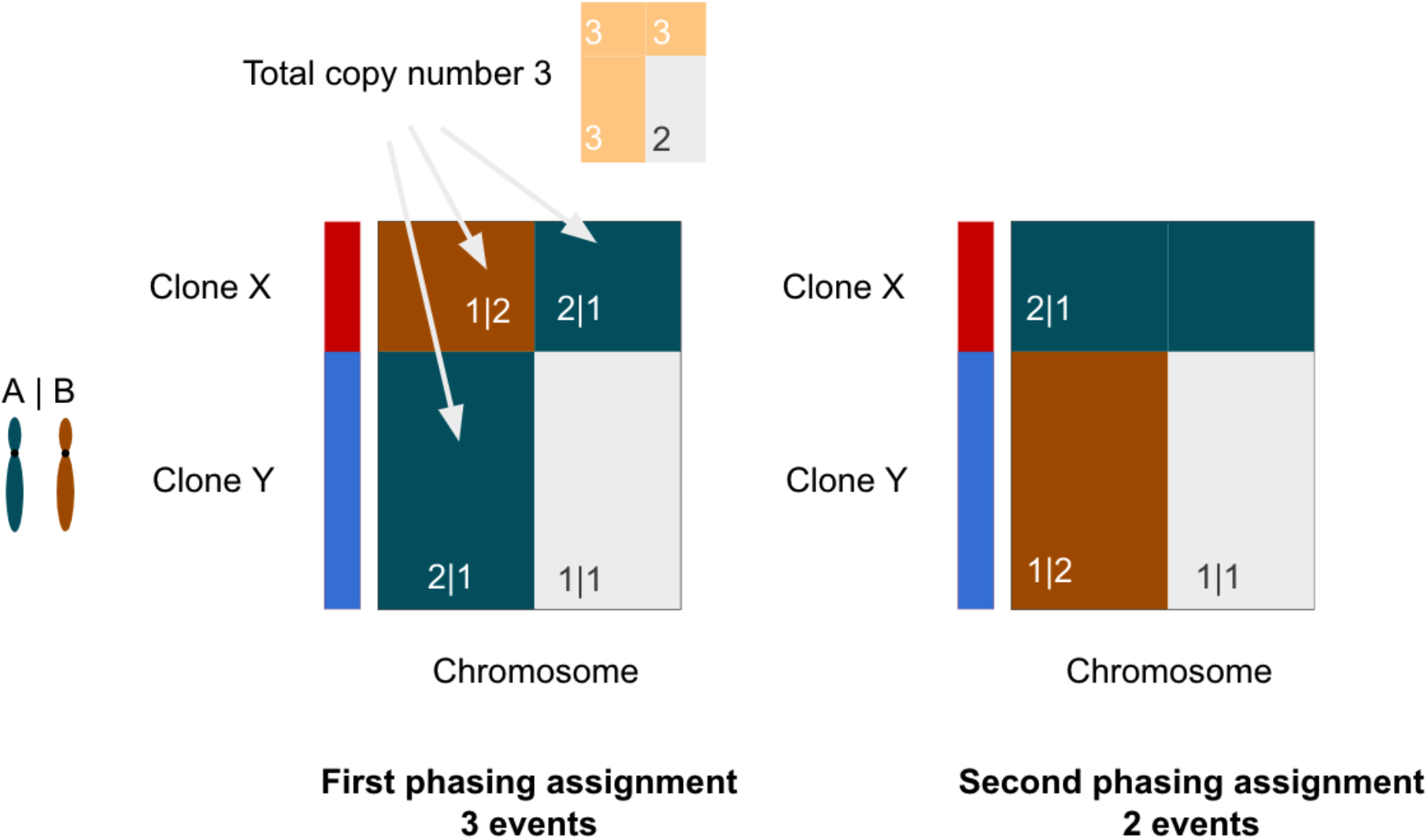
Diagram of rationale behind two-step phasing procedure employed by schnapps. On the left assignment of haplotypes based on the global minimum results in clone X having an event which switches phase half way though the chromosome, the more parsimonious explanation is that this is a single whole chromosome gain and that clone Y has a chromosome arm gain on the opposite haplotype. The second phasing assignment in schnapps attempts to correct for this possibility.

See excel spreadsheet for the following:

**Supplementary Table 1** - Cohort statistics including number of cells per sample, average coverage and number of samples

**Supplementary Table 2** - Table describing the evidence that attributed complex events to breakage fusion bridge cycles

**Supplementary Table 3** - fraction of genome altered as a function of CCF for all samples

**Supplementary Table 4** - chromosome event rates per sample

**Supplementary Table 5** - genomic coordinates of parallel copy number events

## References

1. van Jaarsveld, R. H. & Kops, G. J. P. L. Difference Makers: Chromosomal Instability versus Aneuploidy in Cancer. Trends Cancer Res. 2, 561–571 (2016).

2. Bielski, C. M. et al. Genome doubling shapes the evolution and prognosis of advanced cancers. Nat. Genet. 50, 1189–1195 (2018).

3. Bolhaqueiro, A. C. F. et al. Ongoing chromosomal instability and karyotype evolution in human colorectal cancer organoids. Nat. Genet. 51, 824–834 (2019).

4. Li, Y. et al. Patterns of somatic structural variation in human cancer genomes. Nature 578, 112–121 (2020).

5. ICGC/TCGA Pan-Cancer Analysis of Whole Genomes Consortium. Pan-cancer analysis of whole genomes. Nature 578, 82–93 (2020).

6. Wang, Y. K. et al. Genomic consequences of aberrant DNA repair mechanisms stratify ovarian cancer histotypes. Nat. Genet. 49, 856–865 (2017).

7. Maley, C. C. et al. Genetic clonal diversity predicts progression to esophageal adenocarcinoma. Nat. Genet. 38, 468–473 (2006).

8. Jamal-Hanjani, M. et al. Tracking the Evolution of Non-Small-Cell Lung Cancer. N. Engl. J. Med. 376, 2109–2121 (2017).

9. Macintyre, G. et al. Copy number signatures and mutational processes in ovarian carcinoma. Nat. Genet. 50, 1262–1270 (2018).

10. Watkins, T. B. K. et al. Pervasive chromosomal instability and karyotype order in tumour evolution. Nature (2020) doi:10.1038/s41586-020-2698-6.

11. Laks, E. et al. Clonal Decomposition and DNA Replication States Defined by Scaled Single-Cell Genome Sequencing. Cell 179, 1207–1221.e22 (2019).

12. Minussi, D. C. et al. Breast tumours maintain a reservoir of subclonal diversity during expansion. Nature 1–7 (2021).

13. Sanders, A. D. et al. Single-cell analysis of structural variations and complex rearrangements with tri-channel processing. Nat. Biotechnol. 1–12 (2019).

14. Zaccaria, S. & Raphael, B. J. Characterizing allele- and haplotype-specific copy numbers in single cells with CHISEL. Nat. Biotechnol. 39, 207–214 (2021).

15. Wu, C.-Y. et al. Integrative single-cell analysis of allele-specific copy number alterations and chromatin accessibility in cancer. Nat. Biotechnol. (2021) doi:10.1038/s41587-021-00911-w.

16. McGranahan, N. et al. Allele-Specific HLA Loss and Immune Escape in Lung Cancer Evolution. Cell 171, 1259–1271.e11 (2017).

17. Umbreit, N. T. et al. Mechanisms generating cancer genome complexity from a single cell division error. Science 368, (2020).

18. Campbell, K. R. et al. clonealign: statistical integration of independent single-cell RNA and DNA sequencing data from human cancers. Genome Biol. 20, 54 (2019).

19. Dorri, F. et al. Efficient Bayesian inference of phylogenetic trees from large scale, low-depth genome-wide single-cell data. bioRxiv 2020.05.06.058180 (2020) doi:10.1101/2020.05.06.058180.

20. Funnell, T., O’Flanagan, C., Williams, M. J., Aparicio, S. & P., S. S. The impact of mutational processes on structural genomic plasticity in cancer cells. Biorxiv (2021).

21. Bolnick, D. I., Barrett, R. D. H., Oke, K. B., Rennison, D. J. & Stuart, Y. E. (Non)Parallel Evolution. Annu. Rev. Ecol. Evol. Syst. 49, 303–330 (2018).

22. Gerlinger, M. et al. Genomic architecture and evolution of clear cell renal cell carcinomas defined by multiregion sequencing. Nature Publishing Group doi:10.1038/ng.2891.

23. McClintock, B. The Stability of Broken Ends of Chromosomes in Zea Mays. Genetics 26, 234–282 (1941).

24. Gisselsson, D. et al. Chromosomal breakage-fusion-bridge events cause genetic intratumor heterogeneity. Proc. Natl. Acad. Sci. U. S. A. 97, 5357–5362 (2000).

25. Hadi, K. et al. Distinct Classes of Complex Structural Variation Uncovered across Thousands of Cancer Genome Graphs. Cell 183, 197–210.e32 (2020).

26. Zakov, S., Kinsella, M. & Bafna, V. An algorithmic approach for breakage-fusion-bridge detection in tumor genomes. Proc. Natl. Acad. Sci. U. S. A. 110, 5546–5551 (2013).

27. Shen, M. M. Chromoplexy: a new category of complex rearrangements in the cancer genome. Cancer cell vol. 23 567–569 (2013).

28. Kim, H. et al. Extrachromosomal DNA is associated with oncogene amplification and poor outcome across multiple cancers. Nat. Genet. 52, 891–897 (2020).

29. Wu, C.-Y. et al. Alleloscope: Integrative single cell analysis of allele-specific copy number alterations and chromatin accessibility in cancer. Cold Spring Harbor Laboratory 2020.10.23.349407 (2020) doi:10.1101/2020.10.23.349407.

30. Elizalde, S., Laughney, A. M. & Bakhoum, S. F. A Markov chain for numerical chromosomal instability in clonally expanding populations. PLoS Comput. Biol. 14, e1006447 (2018).

31. Cross, W. et al. Stabilising selection causes grossly altered but stable karyotypes in metastatic colorectal cancer. bioRxiv 2020.03.26.007138 (2020) doi:10.1101/2020.03.26.007138.

32. Baca, S. C. et al. Punctuated evolution of prostate cancer genomes. Cell 153, 666–677 (2013).

33. Stachler, M. D. et al. Origins of cancer genome complexity revealed by haplotype-resolved genomic analysis of evolution of Barrett’s esophagus to esophageal adenocarcinoma. bioRxiv 2021.03.26.437288 (2021) doi:10.1101/2021.03.26.437288.

34. Salehi, S. et al. Single cell fitness landscapes induced by genetic and pharmacologic perturbations in cancer. Cold Spring Harbor Laboratory 2020.05.08.081349 (2020) doi:10.1101/2020.05.08.081349.

35. Tarone, R. E. Testing the goodness of fit of the binomial distribution. Biometrika 66, 585– 590 (1979).

36. Lai, D. & Shah, S. HMMcopy: copy number prediction with correction for GC and mappability bias for HTS data. R package version 1, (2012).

37. Ding, J. et al. Feature-based classifiers for somatic mutation detection in tumour–normal paired sequencing data. Bioinformatics 28, 167–175 (2011).

38. Saunders, C. T. et al. Strelka: accurate somatic small-variant calling from sequenced tumor–normal sample pairs. Bioinformatics 28, 1811–1817 (2012).

39. Layer, R. M., Chiang, C., Quinlan, A. R. & Hall, I. M. LUMPY: a probabilistic framework for structural variant discovery. Genome Biol. 15, R84 (2014).

40. McPherson, A., Shah, S. & Sahinalp, S. C. deStruct: accurate rearrangement detection using breakpoint specific realignment. bioRxiv (2017).

41. Delaneau, O., Marchini, J. & Zagury, J.-F. A linear complexity phasing method for thousands of genomes. Nat. Methods 9, 179–181 (2011).

42. McPherson, A. W. et al. ReMixT: clone-specific genomic structure estimation in cancer. Genome Biol. 18, 140 (2017).

43. Wingett, S. W. & Andrews, S. FastQ Screen: A tool for multi-genome mapping and quality control. F1000Res. 7, 1338 (2018).

44. Burleigh, A. et al. A co-culture genome-wide RNAi screen with mammary epithelial cells reveals transmembrane signals required for growth and differentiation. Breast Cancer Res. 17, 4 (2015).

45. Butler, A., Hoffman, P., Smibert, P., Papalexi, E. & Satija, R. Integrating single-cell transcriptomic data across different conditions, technologies, and species. Nat. Biotechnol. 36, 411–420 (2018).

46. Stuart, T. et al. Comprehensive Integration of Single-Cell Data. Cell 177, 1888–1902.e21 (2019).

47. Wolock, S. L., Lopez, R. & Klein, A. M. Scrublet: Computational Identification of Cell Doublets in Single-Cell Transcriptomic Data. Cell Syst 8, 281–291.e9 (2019).

48. Huang, X. & Huang, Y. Cellsnp-lite: an efficient tool for genotyping single cells. Bioinformatics (2021) doi:10.1093/bioinformatics/btab358.

49. Alquicira-Hernandez, J. & Powell, J. E. Nebulosa recovers single cell gene expression signals by kernel density estimation. Bioinformatics (2021) doi:10.1093/bioinformatics/btab003.

50. McInnes, L. & Healy, J. Accelerated Hierarchical Density Clustering. arXiv [stat.ML] (2017).

51. McInnes, L., Healy, J. & Melville, J. UMAP: Uniform Manifold Approximation and Projection for Dimension Reduction. arXiv [stat.ML] (2018).

52. Louca, S. & Doebeli, M. Efficient comparative phylogenetics on large trees. Bioinformatics 34, 1053–1055 (2018).

53. Werner, B. et al. Measuring single cell divisions in human tissues from multi-region sequencing data. Nat. Commun. 11, 1035 (2020).

